# Extravillous trophoblast model shows generation of bioequivalent N-glycans can maintain immunological protection against natural killer cell cytotoxicity

**DOI:** 10.64898/2026.08.09.743710

**Authors:** Zhengyuan Huang, Alex T. H. Cocker, Guy S. Whitley, Xiangwei Fu, Mark R. Johnson

**Affiliations:** Department of Metabolism, Digestion and Reproduction, Imperial College London, London, UK; Department of Life Sciences, Imperial College London, SW7 2AZ, London, UK; Vascular Biology Research Centre, Molecular and Clinical Sciences Research Institute, City St George’s, University of London, London, UK; National Engineering Laboratory for Animal Breeding, Key Laboratory of Animal Genetics, Breeding and Reproduction of the MARA, Beijing Key Laboratory for Animal Genetic Improvement, State Key Laboratory of Animal Biotech Breeding, College of Animal Science and Technology, China Agricultural University, Beijing, China

**Keywords:** extravillous trophoblast model, N-glycosylation, cell surface N-glycan remodelling, HLA-G, immunomodulatory effect, cytotoxicity assay

## Abstract

Extravillous trophoblasts (EVTs) are a trophoblast subpopulation critical for feto-maternal tolerance during early pregnancy, primarily using HLA-G to exert immunomodulatory effect, and possessing N-glycomic profiles distinct from other trophoblast subpopulations. However, whether the N-glycosylation confers distinct immunological properties to EVTs remains poorly understood. To investigate this, we employed JEG-3, a human choriocarcinoma cell line having the capacity to produce pregnancy-related hormones and expressing both HLA-C and HLA-G resembling placental EVTs, as an *in vitro* EVT model, alongside cell line JAR which exhibits villous trophoblast phenotypes distinct from JEG-3. Both cell lines were treated with kifunensine or swainsonine, inhibitors of α-mannosidases, to remodel their N-glycosylation patterns. This led to significant remodelling of their N-glycomic profiles, with JEG-3 cells showing an increased level of polylactosamine chains and decreased levels of α-2,6-sialylation and core α-1,6-fucosylation. Western blot analysis showed that inhibiting α-mannosidases altered only the composition of N-glycans on cell-surface HLA-G, without affecting the overall abundance of cell-surface HLA-G. In kifunensine-treated JEG-3 cells that predominantly express oligomannose type N-glycans, an intracellular accumulation of unfolded HLA-G fragments, increased hCG secretion, and down-regulations of EVT markers GATA3 and KRT7 were observed compared to untreated control, while swainsonine treatment did not impact N-glycan expression. Cytotoxicity assays using NK-92 as effector cells showed that the de-sialylation of JEG-3 by neuraminidase treatment led to increased NK-92 mediated killing. JEG-3 cell sustained its EVT immunological properties through generating bioequivalent N-glycans, exemplified by NK-92 cells pre-conditioned with used culture media of kifunensine-treated JEG-3 cells displaying reduced cytotoxicity toward NK-sensitive lymphoblast cell line K562, an effect not observed with swainsonine-treated JEG-3 cells. This model suggests that EVTs immunological properties are dependent on specific N-glycomic profiles that are maintained by unique N-glycosylation homeostasis, and overall improves our understanding of how EVTs maintain their immunomodulatory effect at the maternal-fetal interface.

## 1 Introduction

N-glycosylation is a multistep posttranslational modification initiated in the endoplasmic reticulum (ER). The maturation of N-glycans attaching to polypeptide is achieved by a range of finely tuned processes including de-mannosylation, N-acetylglucosamine (GlcNAc) branching, poly-N-acetyllactosamine (poly-LacNAc) extension, and further modification with fucose, sialic acid, and sulphate residues to form various terminal epitopes along the secretory pathway in the Golgi apparatus (Figure 1A). This complex biosynthesis network results in a wide variety of N-glycan structures attached to glycoprotein with N-glycosylation sites of three major types: oligomannose, hybrid, and complex [1–3]. At the molecular level, N-glycosylation has been related to folding, quality control, stability, and transportation of protein, exhibiting multiple biological functions [3,4]. At the cellular level, by virtue of the application of diverse small molecule inhibitors of mammalian N-glycosylation [5] or knockout of genes related to glycosylation in tumour cell lines, the role of N-glycosylation in tumour immune evasion has been demonstrated [6–9]. Deletion of the gene encoding signal peptide peptidase-like 3 in the B cell lymphoblastoma cell line 721.221 that is sensitive to lysis by Natural Killer (NK) cells, led to an enrichment of N-glycans bearing poly-LacNAc extension, making it more resistant to NK-mediated killing [9]. Inducing peripheral tolerance in the local milieu is a common survival strategy for several tumours that prevents their elimination by the host immune system [10]. Mature N-glycan architectures are altered in many malignancies [11–13], which potentially affect tumour immunity by influencing direct interactions with glycan binding proteins expressed on immunomodulatory cell surface [14].

**Figure 1.**
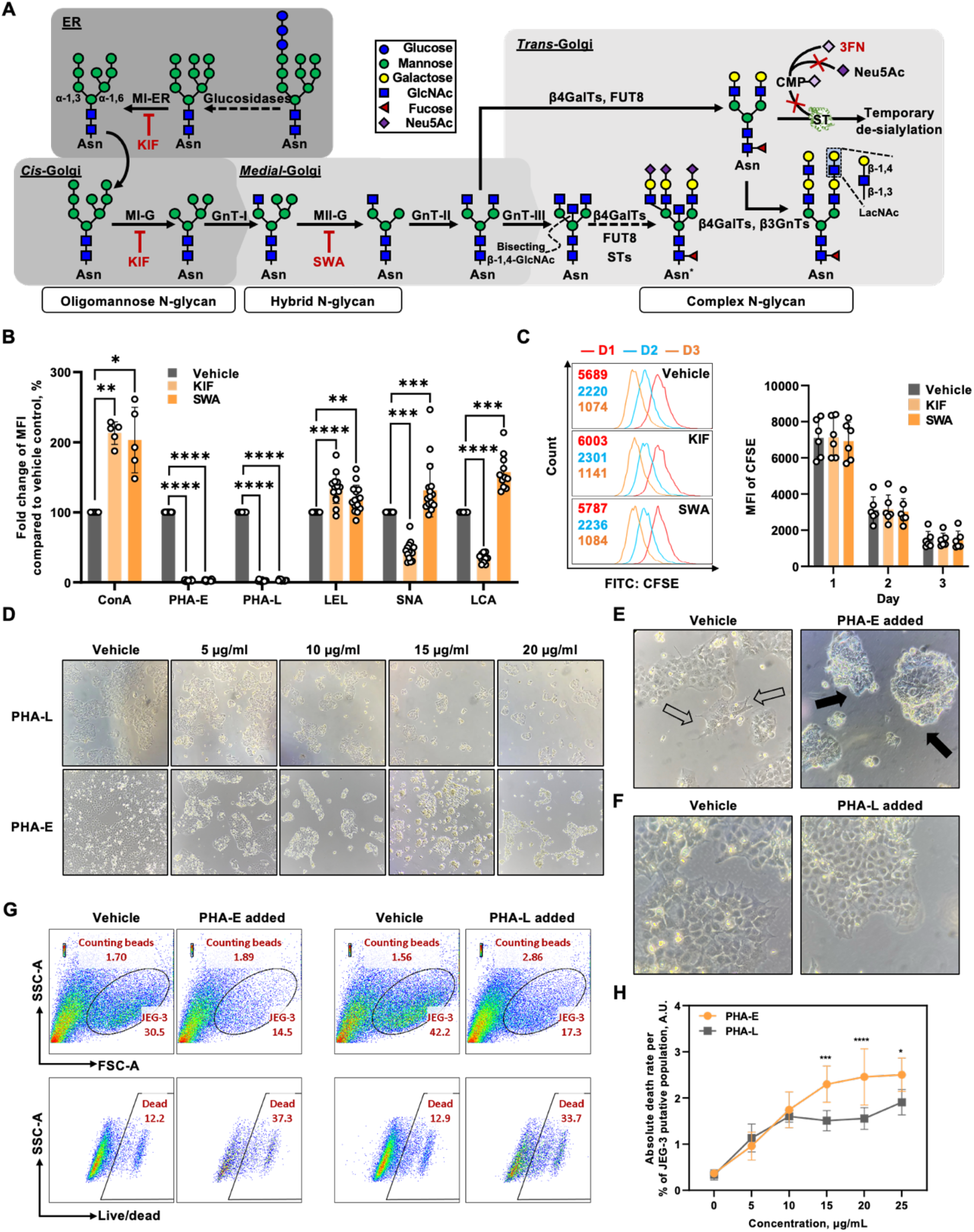
JEG-3 cells are able to generate bioequivalent glycans after the inhibition of α-mannosidases but are susceptible to the masking of functional N-glycan structures. (A) Simplified N-glycosylation pathway in mammalian cells, highlighting the sites of KIF, SWA, and 3FN action and the representative structures expected. The asterisk highlights a complex bisected N-glycan which is found on primary villous cytotrophoblasts [64]. β3GnT, β-1,3-N-acetylglucosaminyltransferase; β4GalT, β-1,4-galactosyltransferase; CMP, cytidine monophosphate; FUT8, α-1,6-fucosyltransferase; GnT, N-acetylglucosaminyltransferase; MI-ER/-G, α-mannosidase I resident in ER/Golgi apparatus; MII-G, α-mannosidase II resident in Golgi apparatus; Neu5Ac, N-acetylneuraminic acid; ST, sialyltransferases. (B) Flow cytometry of vehicle control, KIF-treated, and SWA-treated JEG-3 cells using staining of representative lectins to detect the glycophenotype of the cell surface. Data of KIF- and SWA-treated JEG-3 cells were presented in percentage of change compared with vehicle controls. Same as below. N = 5-12. Paired Student’s t-test was used to compare the average MFI between vehicle control and either of inhibitor-treated JEG-3 cells. (C) Flow cytometry of vehicle control, KIF-treated, and SWA-treated JEG-3 cells to compare the rate of cell proliferation by determining the CFSE fading over three-day incubation. N = 6. Paired Student’s t-test was used to compare the MFI means of CFSE between vehicle control and either of inhibitor-treated JEG-3 cells at each timepoint. (D) The confluency of JEG-3 cell monolayer after treatment with PHA-E or -L in different doses for 48 hours. Images are shown at 40 × magnification. The morphological features of JEG-3 cell in the absence and presence of lectin (E) PHA-E or (F) PHA-L. The viability of JEG-3 cell was determined by flow cytometry after 2-day culture in the presence of either PHA-E or -L. (G) Cells in each treatment were harvested after 48 hours, and then rinsed, stained with live/dead stain, combined with 10 µl of counting beads solution (∼1000 beads/µl) and analysed by flow cytometry. Representative plots are shown. SSC-A against SSC-H plot is first used to gate in singlets and counting beads, afterwards SSC-A against FSC-A plot was used to determine the size and complexity of JEG-3 cell (the shape of gate was based on the vehicle control), and finally the dead population was plotted. FSC-A, forward scatter - area; SSC-A, side scatter - area. (H) Death rate per unit between PHA-E and -L treatments in different concentrations. The ratio of absolute dead rate to the proportion of JEG-3 cell in SSC-A-FSC-A plot is defined as the “death rate per unit”. Data were compared using two-way ANOVA with Bonferroni’s post-hoc test. N = 8 for each data point. A.U., arbitrary units.

Extravillous trophoblasts (EVTs) are originated from cytotrophoblasts in the tips of anchoring villi, and those which invade into decidual stroma termed interstitial cytotrophoblasts, provoking numerous effects during early pregnancy through interacting with decidual stromal and immune cells, eventually exhibiting immunomodulatory effects at a unique tolerized immune compartment called maternal-fetal interface to support the placental development [15–18]. How EVTs fail to elicit immunological rejection in an immune-competent pregnancy despite expressing allogeneic antigens and interacting extensively with decidual immune cells [19] has been the focus of extensive research. Although whether EVTs acquiring immunosuppressive activities in a similar way as malignant cells remains elusive, abundant evidence have suggested that the N-glycosylation is involved in EVTs orchestrating their extensive but controlled infiltration into decidua and interaction with decidual immune cells [20]. Pathological placentae possessing an altered N-glycomic profile compared to that of their healthy counterparts have been observed [21–28], and abnormality in GlcNAc branching of N-glycan has been linked to impaired EVT migration and invasion [29–34]. N-glycosylated proteins expressed on EVT cell surface, such as human leukocyte antigen C (HLA-C) and HLA-G [35,36], interact with receptors on decidual NK cells and macrophages. These interactions lead to altered decidual NK cell cytotoxicity, promotion of regulatory T cell, differentiation of M2 macrophage, and production of anti-inflammatory or growth-promoting cytokines [37–41]. The N-glycosylation of HLA-C has also been related to its cellular localization, correct folding in ER with the involvement of lectin-like chaperones calnexin and calreticulin, and ligation with NK receptors [42–45]. A range of secreted N-glycosylated proteins exerting immunomodulatory effects, including human chorionic gonadotropin (hCG), pregnancy-specific glycoproteins, soluble PD-L, have also been reported to be produced by diverse trophoblast cell lines and primary EVTs [46–55]. In the choriocarcinoma cell line BeWo, which serves as a model of the placental cytotrophoblast layer, pre-treatment with tunicamycin to completely disrupt N-glycosylation before co-culture with peripheral blood-derived NK cells markedly decreases NK cell production of interferon-γ [56]. Knockout and transgenic mouse models demonstrated that the B cell suppression mediated by trophoblast-derived antigens decorated with sialylated N-glycans contributes to the establishment of feto-maternal tolerance, and the absence of sialylated N-glycans leads to embryonic lethality because of a strong alternative pathway-mediated complement activation toward trophoblasts [57,58]. We therefore hypothesised that the glycomic remodelling of EVT proteins can lead to changes in its immunological properties.

This study is the first attempt to investigate the immunological properties of EVT from the glycobiological perspective. We established a simplified model to investigate if the N-glycosylation involved in shaping the immunological properties of EVT. Using the EVT-like choriocarcinoma cell line JEG-3, we found that the N-glycosylation does play a role in regulating and shaping immunological properties of JEG-3 cell. Our results indicate that JEG-3 cells have preferences in expressing poly-LacNAc on cell surface, with immunological properties dependent on specific N-glycomic profiles, and our simplified model has the potential to unveil the link between N-glycomic profile and EVT activities during placentation, providing a new strategy to understanding the altered N-glycomic profile possessed by pathological placentae.

## 2 Materials and methods

### 2.1 Chemicals

All chemicals were purchased from Sigma-Aldrich and media from Thermo Fisher Scientific, unless otherwise indicated. All cell lines were purchased from the American Type Culture Collection and maintained at 37°C and 5% CO_2_ in a humid incubator. All cell cultures were passaged at 80% confluency or at 1×10^6^ cells/ml. Cell passages between 10-30 were used in this study.

### 2.2 Cell culture

Choriocarcinoma cell lines JEG-3 (HTB-36) and JAR (HTB-144), used as *in vitro* models of EVT and villous cytotrophoblast, were respectively incubated in Minimum Essential Medium (MEM) supplemented with 10% (v/v) fetal bovine serum (FBS), 1 mM sodium pyruvate, 1× non-essential amino acid, 100 units/ml of penicillin, and 100 μg/ml of streptomycin, and complete RPMI-1640 medium (ATCC modification type) supplemented with 10% (v/v) FBS, 100 units/ml of penicillin, and 100 μg/ml of streptomycin. Human immortalised myelogenous leukaemia cell line K562 (CCL-243) and non-adherent immortal NK cell line NK-92 (CRL-2407), used as target and effector in cytotoxicity assays, were respectively incubated in RPMI-1640 medium (GlutaMAX™ supplement) supplemented with 10% (v/v) FBS, 100 units/ml of penicillin, and 100 μg/ml of streptomycin (referring to “R10”), or complete AMEM medium supplemented with 0.2 mM inositol, 0.02 mM folic acid, 200 IU/ml recombinant human IL-2, 12.5% (v/v) horse serum, 12.5% (v/v) FBS, 100 units/ml of penicillin, 100 μg/ml of streptomycin, and 0.1 mM 2-mercaptoethanol (replenished daily).

### 2.3 Inhibition of N-glycosylation and masking of cell-surface complex N-glycan

Cells were cultured in the presence of mannosidase inhibitors (Supplementary Table 1) for 3 d to reduce the presence of hybrid and native complex N-glycans (Figure 1A) on cellular glycoproteins as previously described [59]. The cells incubated with vehicle (dimethyl sulfoxide and/or buffer) were used as controls. As for western blotting analyses of conditioned media, the first conditioned media of treated JEG-3 cell were discarded after 2-day treatment and then fresh supplemented MEM without FBS in the absence or presence of mannosidase inhibitors were replenished. Cells were incubated for another 2 d. Afterwards, the conditioned media were collected, concentrated by 3 kDa molecular weight cutoff Amicon® Ultra Centrifugal Filter (Merck) by following the manufacturer’s instructions, and stored at −20°C for future use. To reduce the presence of sialylated N-glycans on cellular glycoproteins (Figure 1A), JEG-3 cells were cultured in the presence of sialyltransferase inhibitors (Supplementary Table 1) for 3 d as previously described [60]. To reduce the presence of core 2-type O-glycan, JEG-3 cells were cultured in the presence of inhibitor targeting C1Gal-Ts (Supplementary Table 1) for 3 d. JEG-3 cells were also cultured in the presence of lectins *Phaseolus vulgaris* erythronagglutinin or leukoaggulutinin (PHA-E or -L, Supplementary Table 2) at different doses (5, 10, 15, 20, 25 μg/ml) for 2 d to mask cell surface complex N-glycans. Treated cells were analysed by flow cytometry, or lysed and then subjected to either Western blotting or immunoprecipitation.

### 2.4 First trimester chorionic villi collection and protein extraction

First trimester placentae were collected from women in first trimester pregnancy (ranged between 6 to 13 weeks of pregnancy) who opted to undergo surgical termination of pregnancy in Chelsea and Westminster Hospital and West Middlesex University Hospital. Ethical approval was obtained from London-Chelsea Research Ethics Committee (REC reference: 11/LO/0971). Samples were processed within 4 h of collection. Placental tissue was rinsed in ice-cold Dulbecco’s Phosphate-Buffered Saline (DPBS) and finely dissected. Residual blood clots were removed by gentle teasing with forceps and scissors. Tissue homogenate was prepared as previously described [61]. In brief, rinsed tissue was transferred to pre-cooled Precellys™ CK-28R tube filled with ice-cold RIPA lysis buffer and then subjected to homogenization executed by Precellys™ homogenizer (Bertin Technologies SAS). Homogenates were rested on ice in-between cycles. The homogenate was centrifuged at 10,000 × g for 10 min at 4°C; the supernatant was transferred to a new 1.5 ml microfuge tube and stored at −20°C. Protein concentrations were determined using Pierce™ BCA Protein Assay Kits (Thermo Fisher Scientific) by following the manufacturer’s instructions.

### 2.5 Glycomic profiling using lectin staining

Cells were washed in DPBS, resuspended with FACS (fluorescence-activated cell sorting) buffer (formula: DPBS, pH 7.2, 1% FBS, and 2 mM EDTA) supplemented with 10 μg/ml of any fluorophore-conjugated lectins (Vector Laboratories; Supplementary Table 2), and then incubated at 4°C for 15 min. Cells were washed in FACS buffer and combined with 10% (v/v) Precision Count Beads™ (BioLegend) before FACS analysis on a BD LSRFortessa™ cell analyser (BD Biosciences). FACS data analyses were performed using FlowJo™ Software workspace (Tree Star).

### 2.6 Proliferation assays

The cell proliferation over 72-hour treatment was studied using the CellTrace™ CFSE (carboxyfluorescein succinimidyl ester) Cell Proliferation Kit (Invitrogen) according to the manufacturer’s instruction. Cells were suspended in CFSE staining solution, incubated at 37℃ for 20 min, rinsed with complete culture medium, and then resuspended in fresh, pre-warmed complete culture medium in the presence or absence of inhibitors. For each condition, approximately 1-2×10^5^ cells were separately seeded into three wells of 24-well plate, allowing the CFSE intensity to be analysed at 24, 48 and 72 hrs after seeding.

### 2.7 Determination of HLA-G distribution

Two clones (MEM-G/9 and 4H84) of anti-HLA-G monoclonal antibody (mAb) with different characteristics (Supplementary Table 3) were used to study the distribution and folding status of HLA-G expressed by JEG-3 cell. Cells were first stained with clone MEM-G/9 anti-HLA-G (1 in 100 dilution; Abcam), conjugated with either phycoerythrin (PE), fluorescein isothiocyanate (FITC), or allophycocyanin (APC), and then subjected to fixation and/or permeabilization using the BD Cytofix/Cytoperm™ Fixation/Permeabilization Kit (BD Biosciences). Fixed and/or permeabilized cells were stained with clone 4H84 anti-HLA-G (1 in 100 dilution; Santa Cruz Biotechnology) conjugate with Alexa Fluor (AF) 488 and subjected to flow cytometric analyses.

### 2.8 SDS-PAGE and Western blotting

All equipment and chemicals used in electrophoresis and Western blotting were purchased from Bio-Rad unless otherwise indicated. Treated cells were lysed for 30 min in ice-cold RIPA, of which protein concentration was determined by bicinchoninic acid assay. The cell lysis was combined with Laemmli sample buffer supplemented with 50 mM dithiothreitol and denatured at 96°C for 6 min. For each condition, 20 μg of denatured protein samples were loaded and subjected to separation by SDS-PAGE under reducing conditions and then proteins were transferred to an PVDF membrane. Membranes were probed with either mouse anti-human HLA-G (clone 4H84), mouse anti-human GAPDH (clone 6C5; both from Santa Cruz Biotechnology), mouse anti-human β-actin (clone 8H10D10, Cell Signaling Technology) mAbs, or rabbit polyclonal anti-human hCG (Abcam), followed by HRP (horseradish peroxidase)-conjugated secondary antibodies (Cell Signaling Technology). Visualization was performed using Clarity Western ECL Substrate and iBright™ FL1500 imaging system (Thermo Fisher Scientific). Data capture and analysis was conducted using iBright™ Analysis Software (Thermo Fisher Scientific) for all membranes.

### 2.9 Protein de-glycosylation by glycosidase digestion

Whole cell lysates of JEG-3 cells and protein extracts of first trimester placenta were combined with denaturing buffer and heated at 96°C for 6 min. Denatured samples are further incubated in reaction buffer containing endoglycosidase (Endo) H or Peptide-N-Glycosidase (PNGase) F (New England BioLabs). After 1 h incubation at 37℃, glycosidase-digested samples were subjected to SDS-PAGE and Western blotting.

### 2.10 Immunoprecipitation of HLA-G molecules

All equipment and chemicals used in immunoprecipitation were purchased from Invitrogen unless otherwise stated. Dynabeads^®^ protein A/G coupled with mouse anti-HLA-G mAb (clone G233, Abcam; Supplementary Table 3) was incubated with whole cell lysate of JEG-3 cell at 4°C overnight on a tube rotator. The DynaMag™-2 magnet pelleted the HLA-G-Dynabeads^®^ protein A/G complex. Post-immunoprecipitation supernatant was collected and stored at −20°C for further use. The remaining pellet was rinsed and resuspended in 80 μl of Laemmli sample buffer (Bio-Rad) complemented with 50 mM dithiothreitol, mixed well, heated at 96°C for 6 min, and cooled down to room temperature before placing back on the DynaMag™-2 magnet. Supernatant was collected upon beads forming a tight pellet and stored at −20°C for further use.

### 2.11 ELISA for hCG

The levels of whole hCG secreted by JEG-3 cell into culture media were determined using Human hCG ELISA Kit (Abcam) according to the manufacturer’s instruction. In brief, whole hCG molecules in standards and diluted supernatant samples were first captured by anti-hCGα mAbs immobilized on a 96-well microplate; afterwards, anchored molecules were detected by biotinylated anti-human hCGβ mAb and then HRP-conjugated streptavidin was used to quantify the level of whole hCG in each well based on colorimetric analysis. Microplate Manager Software 6 (Bio-Rad) was used to generate a graph for the standard curve (450 nm absorbance versus hCG standard concentration) and then hCG concentration of each sample was determined based on the standard curve. All absorbance values for standards and samples were corrected using those from the zero standard.

### 2.12 Flow cytometry

The following anti-human antibodies were used for surface staining: anti-CD56 Brilliant Violet 650 (BV650; clone NCAM16.2, BD Biosciences), anti-HLA-G FITC (clone MEM-G/9, Abcam). For intracellular staining, the following antibodies were employed: anti-GATA3 Brilliant Blue 700 (clone L50-823, BD Biosciences), anti-KRT7 AF647 (clone W16155A, BioLegend), anti-IL-8 PE (clone E8N1, BioLegend), anti-IFN-γ AF700 (clone 4S.B3, BioLegend), and anti-CD107a PE-Cyanine7 (clone H4A3, BioLegend).

Cells were first stained with a fixable viability dye (Live/Dead Fixable Aqua Dead Cell Stain Kit, Life Technologies) together with the appropriate surface antibodies for 15 min at 4°C. For intracellular staining, cells were fixed and permeabilized using the BD Cytofix/Cytoperm™ Fixation/Permeabilization Kit according to the manufacturer’s instructions, followed by incubation with intracellular antibodies for 30 min at 4°C. Excess antibodies were removed by washing the cells (500 × g, 5 min, 4°C) after each staining step and twice following the final intracellular antibody incubation. Flow cytometry-based cytotoxicity assay was set up as previously described [62] with modification (Supplementary Figure 3A). In general, either effectors or targets were pre-treated and mixed to reach an effector-to-target ratio at 5:1, and then co-incubated at 37°C for 10 h. In parallel, targets were cultured under the same condition in the absence of effector to determine their spontaneous death rate. All cultures were performed in at least triplicate. For verifying the effect of de-sialylation on JEG-3 cell immunogenicity, adherent JEG-3 cell monolayer was treated with 200 IU/ml of α2-3,6,8 neuraminidase (New England Biolabs) for 1 h at 37°C, gently rinsed with pre-warmed DPBS twice afterwards, and then co-incubated with five-fold of NK-92 cells in fresh R10 supplemented with 200 IU/ml IL-2 (referring to “R10/IL-2” hereafter) for 10 h. After treatment, supernatant and inhibitor-treated JEG-3 cell were collected separately to verify the effect of glycan remodelling on immunoregulatory effects of JEG-3 cell. Before the co-culture with K562 cell, NK-92 cell underwent 6-hour pre-conditioning with: (a) equal number of different inhibitor-treated JEG-3 cells in fresh R10/IL-2, (b) supernatant of JEG-3 cell culture after three-day drug treatment, or (c) culture media supplemented with different drugs and calibrated for 3 d in incubator (Supplementary Figure 3A). K562 cells were labelled with CFSE (incubation with 1 μM CFSE in DPBS for 15 min at 37℃) before adding into the well containing NK-92. At the end of co-culture, cells were washed and resuspended in FACS staining buffer, and then successively stained with LIVE/DEAD™ Fixable Near-IR Stain (L/D, dilution 1 in 1000), anti-CD56 BV650 mAb (dilution 1 in 100), and anti-HLA-G FITC mAb (dilution 1 in 100), and eventually subjected to viability assessment by flow cytometry. For the gating strategy (Supplementary Figure 3A), debris and doublets were first excluded, and then targets were identified by gating on HLA-G^+^ CD56^−^ for JEG-3 cell or CFSE^+^ CD56^−^ for K562 cells. L/D positive population was identified, of which the percentage to the total HLA-G^+^ CD56^−^ or CFSE^+^ CD56^−^ population represents the absolute death rate achieved for each condition. Cytotoxicity activity of effector against target was evaluated as previously reported [63] with modification:

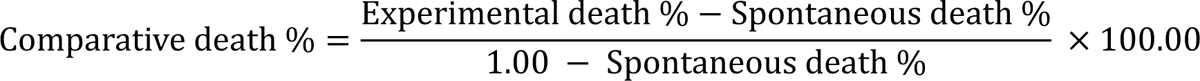

The comparative death rate of K562 cell was the final readout indicating the cytotoxicity of NK-92 cell.

### 2.13 NK-92 cell stimulation during the co-culture with JEG-3 cell

Four hours before the end of co-culture, a mixture comprised of anti-CD107a mAb, cell stimulation cocktail (containing 81 nM phorbol 12-myristate 13-acetate and 1.34 μM ionomycin), 2 μM monensin, and 3.0 μg/ml Brefeldin A (all from Invitrogen), was added into the “Co-culture” well for NK-92 stimulation, while only anti-CD107a mAb, monensin and Brefeldin A were included in the “Co-culture” well without stimulation. After 10-hour co-culture with de-sialylated JEG-3 cells, NK-92 cells were harvested, rinsed and subjected to cell surface and intracellular staining by following manufacturer’s instructions provided for fluorophore-conjugated antibodies. Cells were analysed using flow cytometry.

### 2.14 Quantification and Statistical Analysis

Analysis was performed with Prism version 9.0.0 (GraphPad Software). All data were tested for normality using a Kolmogorov–Smirnoff test or a D’Agostino & Pearson normality test depending on the group size. For normally distributed data, either paired or unpaired two-tailed Student’s t-test was used to perform comparison between two groups, while ANOVA followed by a Dunnett’s or Bonferroni’s post-hoc test for three groups or more. As for non-normally distributed data, either Wilcoxon signed-rank test or Mann-Whitney U test was used to perform comparison between two paired or unpaired groups, while Friedman’s test with a Dunn’s multiple comparisons post-hoc test for three groups or more. Data are shown as mean ± SD based on at least three independently repeated experiments. A *p*-value of less than 0.05 was statistically significant (\**p* < 0.05), \*\**p* < 0.01, \*\*\**p* < 0.001, and \*\*\*\**p* < 0.0001.

## 3 Results

### 3.1 JEG-3 cell is tolerative to pharmacological inhibition of N-glycan processing but not to the lectin masking of cell surface complex N-Glycans

It is well known that the structural diversity of N-glycans requires a series of biosynthesis in tight control [2], but the necessity of possessing N-glycan in diversity for EVT remains poorly understood. We therefore established a simplified model by culturing JEG-3 cells in the presence and absence of the α-mannosidase I inhibitors kifunensine (KIF; 10 μg/ml) or the α-mannosidase II inhibitor swansonine (SWA; 10 μg/ml) for 3 d to reduce the presence of hybrid and native complex N-glycans on cellular glycoproteins (Figure 1A). Cell surface staining with a panel of fluorophore-conjugated lectins (Supplementary Table 2) combined with flow cytometry was applied to determine the level alteration of specific glycan component after treatment. Both KIF and SWA treatment induced increased levels of terminal α-mannose (indicated by ConA binding) and poly-LacNAc (LEL) and complete losses of both β-1,4- and β-1,6-GlcNAc branches (PHA-E and -L), while levels of α-2,6-sialylation (SNA) and core α-1,6-fucosylation (LCA) were differentially affected, with KIF reducing and SWA increasing (Figure 1B). The level of α-2,3-sialylation (MAL II) was not studied because no binding was observed when JEG-3 cells incubated under any doses of fluorophore-conjugated MAL II (Supplementary Figure 1A). Given that KIF and SWA affect all N-glycosylated proteins in the cell, we used identically treated JEG-3 cells in flow cytometry experiments to evaluate cell proliferation over three-day treatment. No significant difference in CFSE intensity being observed between control and either inhibitor-treated JEG-3 cells (Figure 1C). Lectin blot analyses of total cell lysates of inhibitor-treated JEG-3 cells showed a similar pattern of glycomic alteration as the flow cytometric analyses but with more details about what molecular weight of glycoprotein the glycomic alteration is restricted to (Supplementary Figure 1B). Although the inhibitor-induced absences of cell surface complex N-glycan (indicated by the absence of PHA-E or -L binding) exerted no detrimental effect on JEG-3 cell proliferation, we found that masking the N-glycans expressed on cell surface by either PHA-E or -L markedly restricted JEG-3 cell proliferation (Figure 1D); notably, JEG-3 cells cultured in the presence of PHA-E underwent morphological alterations characterised by the formation of cell spheres without any pseudopodium-like projections (solid arrows) compared with the control (hollow arrows, Figure 1E), while those of PHA-L remained adherent as the control (Figure 1F). Flow cytometric analyses further confirmed that both lectins induced JEG-3 cell death which was more prominent with PHA-E added (Figures 1G, H). Overall, we found that losing the ability of synthesizing and presenting complex N-glycan on cell surface did not impair the proliferation of JEG-3 cell, whereas masking these complex N-glycans markedly reduced their proliferative capacity.

### 3.2 JEG-3 cell is prone to generate bioequivalent glycans to maintain a constant level of poly-LacNAc

JEG-3 cell is not only unaffected by the pharmacological inhibition of N-glycan processing in term of proliferation, but also capable of generating bioequivalent glycans in response to this inhibition. Comparisons in N-glycomic profiles among three primary trophoblast subtypes have concluded that the N-glycomic profile of EVT is different from those of syncytiotrophoblast and cytotrophoblast [64]. To determine whether the observe tolerance and capacity are restricted to EVT, we also cultured JAR (as villous cytotrophoblast cell model) and K562 (as non-trophoblast cell model) cells in the presence and absence of KIF or SWA for 3 d as JEG-3 cell. We found that JAR cells shared a similar pattern of glycomic alteration with JEG-3 cells after both treatments, with the exception of no significant difference in the level of α-2,6-sialylation between SWA-treated samples and control (Figure 2A), while K562 cells exhibited differences in the pattern of glycomic alteration when compared to JEG-3 cell, characterized by no significant increase in poly-LacNAc after either KIF or SWA treatments and no increase in α-2,6-sialylation after SWA treatments (Figure 2B). The variation in responses towards pharmacological inhibition of N-glycan processing between cell lines indicated they were appropriate models to explore the regulatory role of N-glycosylation in EVTs. Although both JEG-3 and JAR cells exhibited an increased level of poly-LacNAc after either KIF or SWA treatment, we noticed that the former had greater increases (Figure 2C) based on the comparison in percentage change of LEL binding relative to control. In fact, the KIF treatment theoretically results in the absence of potential GlcNAc branches that can be elongated by poly-LacNAc, whereas an increased LEL binding was still observed in KIF-treated JEG-3 cell (Figure 1B). Considering that poly-LacNAc is a glycan component shared by both N- and O-glycans [65,66], we therefore questioned if this increase is a consequence of abundant poly-LacNAc extensions decorating core 2-type O-glycan. We treated JEG-3 cells with benzyl-α-GalNAc (BAG, Supplementary Figure 1), an inhibitor of the biosynthesis of core 2-type O-glycan precursor without inhibitory activity on polypeptide N-acetyl-α-galactosaminyltransferases or N-linked glycoprotein secretion [5,67], in the absence or presence of KIF. Instead of decreasing, the co-treatment of KIF and BAG even induced a greater increase in level of poly-LacNAc when compared to KIF only (Figure 2D). An increase in level of bisecting β-1,4-GlcNAc was also observed in BAG-treated cell (Figure 2D). These results collectively demonstrate that the compensatory increase of poly-LacNAc after pharmacological inhibition of N-glycan processing is restricted to cell lines with trophoblastic origin, while JEG-3 cell is in greater need for a constant level of poly-LacNAc than JAR cell.

**Figure 2.**
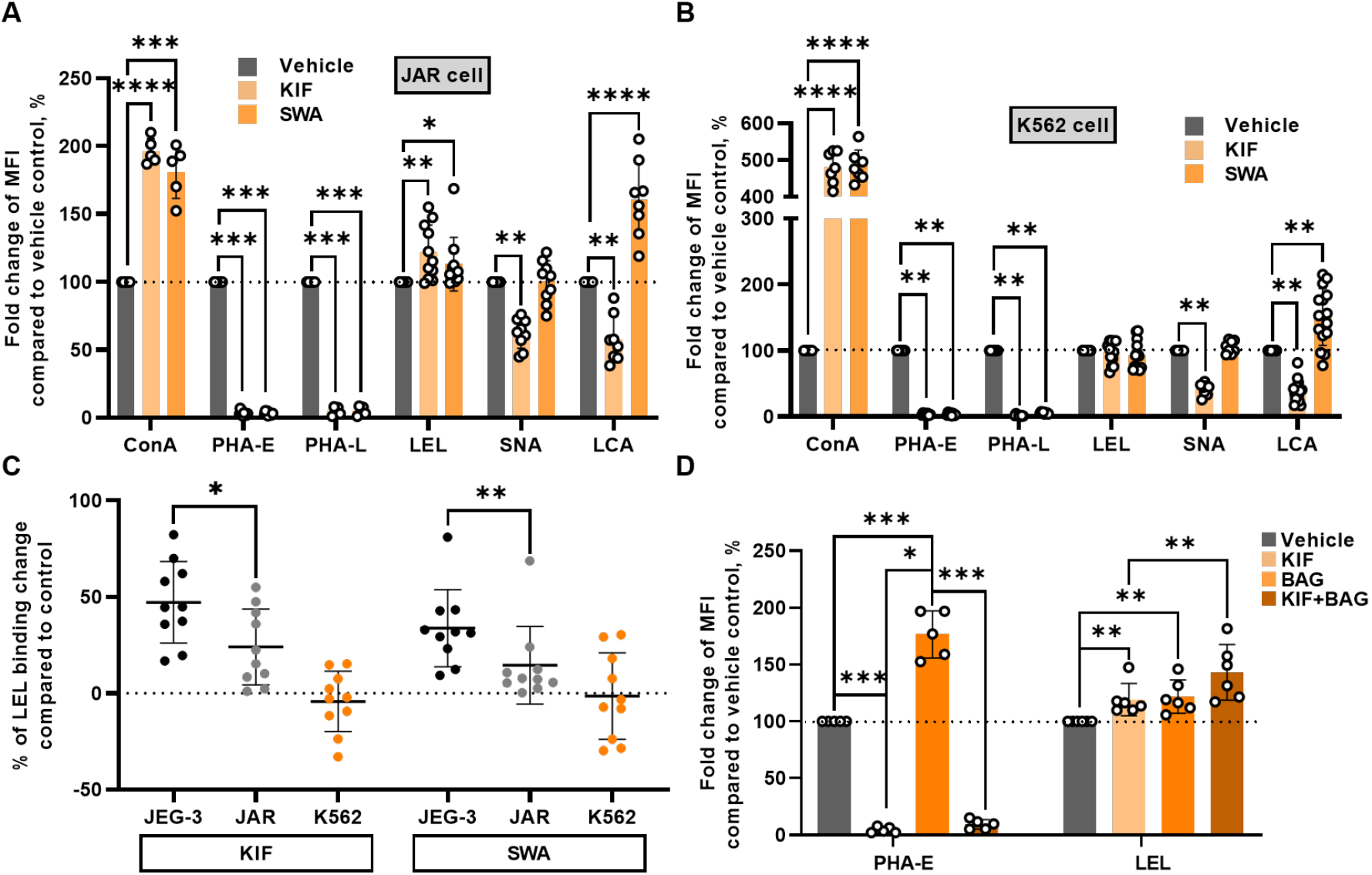
JEG-3 cell has preference in generating excessive poly-LacNAc extension in response to the inhibition of glycosylation. Flow cytometry of **(A)** inhibitor-treated JAR cells and their vehicle control (N = 5-11), and **(B)** inhibitor-treated K562 cells and their vehicle control (N = 7-15), using staining of representative lectins to detect the glycophenotype of the cell surface. Paired Student’s t-test was used to compare the average MFI between vehicle control and either of inhibitor-treated cells. **(C)** The extent of MFI change of LEL staining in inhibitor-treated cell relative to that in vehicle control. Either unpaired Student’s t-test or Mann-Whitney U test was used to compare the means between two cell lines under same treatment and exhibiting the same trend. N = 10. **(D)** PHA-E or LEL staining of JEG-3 cells with KIF and/or BAG treatment, and vehicle control. Paired Student’s t-test was used to compare the average MFI between every two paired treatments. N = 5 or 6.

### 3.3 The composition of attached N-glycans does not affect the cellular localisation and expression of HLA-G1 in JEG-3 cell

JEG-3 cells can maintain a constant level of poly-LacNAc even after the inhibition of both N-and O-glycosylation, we therefore sought to confirm whether they can maintain their glycoprotein expression regardless of glycomic alteration. We first studied the expression of HLA-G which is a well-known EVT-derived glycoprotein possessing one N-glycosylation site and expressing in both membrane-bound and soluble form [36,68–70]. We first confirmed that JEG-3 cell is expressing the HLA-G in large quantities as EVTs under physiological status as evidenced by a 38-kDa band was blotted by the antibody clone 4H84 in both JEG-3 cell lysate and lysates of human first trimester chorionic villi under reducing conditions in Western blotting (Figure 3A, B). However, we also observed that another band with a molecular weight of ∼30 kDa was also blotted in JEG-3 cell lysate (Figure 3A). All these bands were further found to be N-glycosylated as evidenced by the shifts observed for both bands after the PNGase F digestion (Figure 3A, B); moreover, the ∼30-kDa band was found to have a higher signal intensity in KIF-treated JEG-3 cell compared to control (red chevron, Figure 3A). No significant differences in normalized protein abundance of 38-kDa band between KIF-treated JEG-3 cells and control were observed, whereas abundance of the ∼30-kDa band was significantly increased in KIF-treated JEG-3. The abundance of neither 38-kDa nor ∼30-kDa band was affected by the SWA treatment (Figure 3C). The flow cytometry of intact JEG-3 cell using two different clones of anti-HLA-G mAbs (Supplementary Table 3) specifically binding correctly-folded HLA-G and unfolded HLA-G respectively, further confirmed that the 38-kDa band indicates membrane-bound HLA-G1, an HLA-G isoform encoded by a full-length mRNA [71], while the ∼30-kDa band indicates unfolded HLA-G fragment located intracellularly (Supplementary Figure 2A-E). Similar to results from Western blotting, flow cytometric analyses of intact JEG-3 cells confirmed a significant higher proportion of 4H84 positive population in KIF-treated JEG-3 cell when compared to that in control (Figures 3D, E), suggesting an intracellular accumulation of unfolded HLA-G fragment. Dysfunction of N-glycan processing may disrupt the transfer of monosaccharides to one (hybrid type) or both (oligomannose type) branches, making N-glycans sensitive to Endo H digestion [72]. In fact, for both first-trimester chorionic villi and JEG-3 cells, their membrane-bound HLA-G1 are not necessarily decorated with complex N-glycans because there exist N-glycans on membrane-bound HLA-G1 are sensitive to Endo H digestion (yellow and white chevrons, Figure 3B, F), while those unfolded HLA-G fragments with nascent N-glycans located intracellularly were all Endo H-sensitive (red chevron, Figure 3F). When it comes to either KIF or SWA treatment, both membrane-bound HLA-G1 and unfolded HLA-G fragment were sensitive to Endo H digestion (white and red chevrons, Figure 3F), indicating the expression cell-surface HLA-G1 decorated with either oligomannose- or hybrid-type N-glycans. Lectin blotting analyses of HLA-G1 immunoprecipitated from whole cell lysate of inhibitor-treated JEG-3 cell also confirmed that no PHA-E binding was observed for blots in both KIF- and SWA-treated JEG-3 cells (Figure 3G). These results directly demonstrate that for glycoproteins HLA-G, the composition of the attached N-glycans does not affect its proper folding or cell-surface expression.

**Figure 3.**
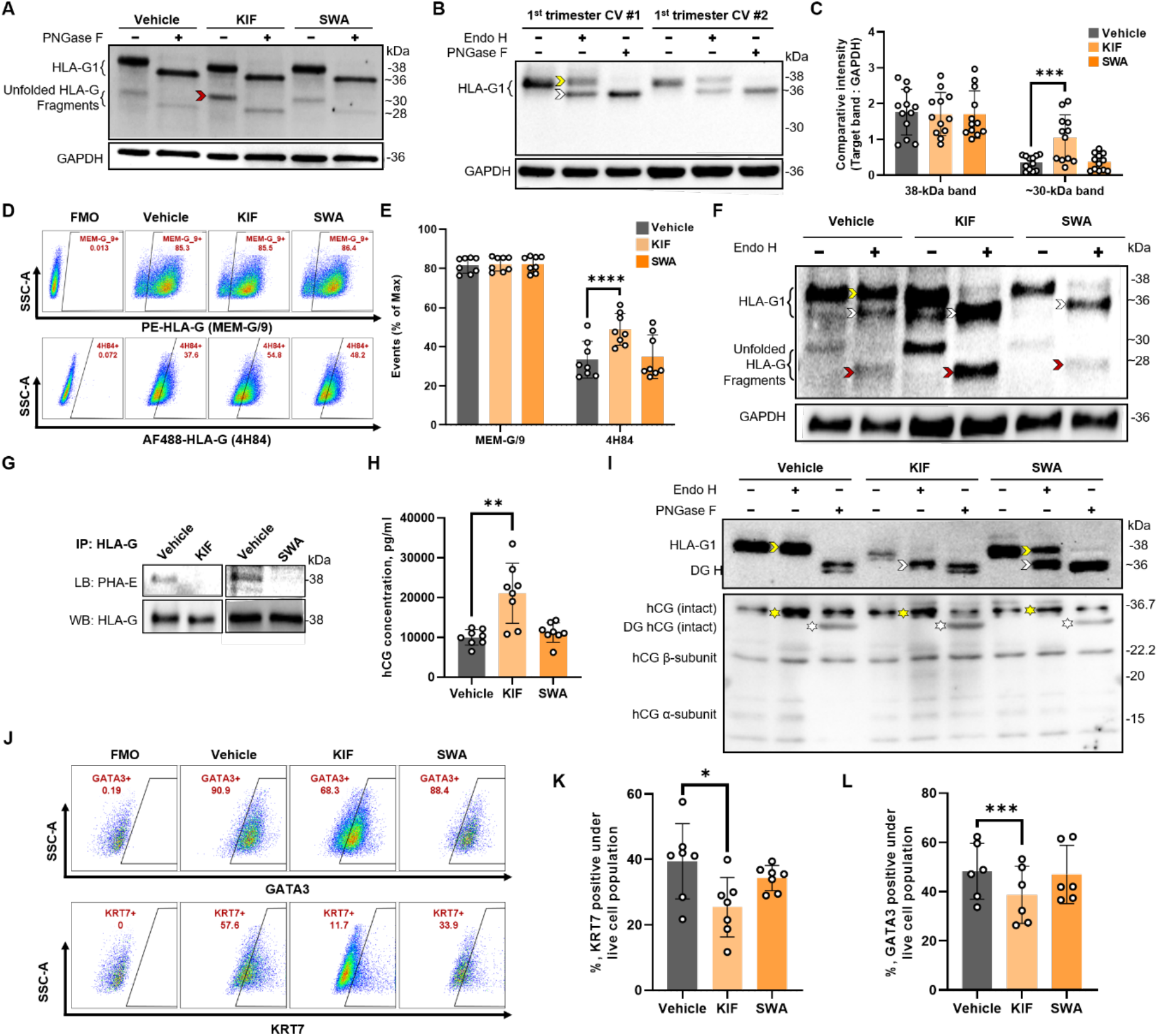
The predominance of oligomannose type N-glycan induces phenotypic abnormalities of JEG-3 cell. **(A)** Both HLA-G1 and unfolded HLA-G fragment are N-glycosylated. Western blots of HLA-G in total lysates of inhibitor-treated JEG-3 cells and vehicle control before and after de-glycosylation by PNGase F treatment. Western blot of GAPDH was used as a loading control. Same as below. Representative blots are shown. **(B)** Western blot analysis of protein extract of first trimester chorionic villi with (+) or without (−) Endo H or PNGase F digestion. **(C)** The comparison in normalized optical density of either HLA-G1 or unfolded HLA-G fragment between inhibitor-treated JEG-3 cell populations and vehicle control. **(D)** Detection of HLA-G on the cell surface of or inside JEG-3 cells by flow cytometric analysis. SSC-A against FSC-A plot was first used to determine the size and complexity of JEG-3 cells. After the exclusion of doublets and dead cells, either MEM-G/9 or 4H84 positive population was plotted. **(E)** The comparison in proportion of MEM-G/9 or 4H84 positive population between inhibitor-treated JEG-3 cell populations and vehicle control. **(F)** Western blot analysis of HLA-G in whole cell lysates of control, KIF-treated, and SWA-treated JEG-3 cells with (+) or without (−) Endo H digestion. **(G)** Lectin blot analysis of PHA-E on immunoprecipitated HLA-G1 from either inhibitor-treated JEG-3 cells or vehicle control. Representative blots are shown. **(H)** The culture media after inhibitor treatment were analyzed for their whole hCG content by ELISA. **(I)** Western blot analysis of soluble HLA-G and hCG from conditioned media of inhibitor-treated JEG-3 cell and vehicle control, with (+) or without (−) Endo H or PNGase F digestion. **(J)** The comparison in normalized optical density of either HLA-G1 or unfolded HLA-G fragment between inhibitor-treated JEG-3 cell populations and vehicle control. Detection of KRT7 and GATA3 inside JEG-3 cells by flow cytometric analysis after drug-treatments. SSC-A against FSC-A plot was first used to determine the size and complexity of JEG-3 cells. After the exclusion of doublets and dead cells, either GATA3 or KRT7 positive population was plotted. JEG-3 cells stained with L/D staining only were plotted to determine the boundary of gates for target positive populations. The comparison in proportion of **(K)** KRT7 or **(L)** GATA3 positive population between inhibitor-treated JEG-3 cell and vehicle control. Either paired two-tailed Student’s t-test or Wilcoxon signed-rank test was used to compare the means between vehicle control and either of inhibitor-treated JEG-3 cells. N = 6-15. CV, chorionic villi; DG, de-glycosylated; Endo H, endoglycosidase H; IP, immunoprecipitated; LB, lectin blotting; PNGase F, peptide-N-glycosidase F; WB, Western blotting.

### 3.4 The secretion of hCG and soluble HLA-G1 is independent of the composition of their attached N-glycans

Apart from membrane-bound glycoprotein HLA-G, we questioned if the glycoprotein secretion by JEG-3 cell is also affected by the inhibition of N-glycan processing. The hCG is a potent immunomodulatory hormone and a high proportion of N-glycans has been determined in the composition of this secreted glycoprotein [73–75]. The concentration of whole hCG molecule released into the conditioned medium from JEG-3 cell with or without inhibitor treatment was studied, and we found that only KIF treatment induced an increased concentration of hCG in conditioned medium when compared to control (Figure 3H). Soluble form of HLA-G1 has also been reported to protect HLA class I-negative K562 cells from NK lysis, acting as a potential immunoregulator in NK cell recognition [76]. We therefore further studied the N-glycans on soluble isoforms of HLA-G1 and hCG secreted by JEG-3 cells through Western blotting analyses of concentrated conditioned media. Similar to the findings with membrane-bound HLA-G1 bands in lysates, soluble HLA-G1 molecules from the conditioned medium migrated faster in KIF- and SWA-treated samples and were Endo H sensitive (white chevrons, Figure 3I), while those in control were all Endo H resistant (yellow chevrons, Figure 3I), suggesting that JEG-3 cell under inhibition of N-glycan processing could still secret soluble HLA-G1 decorated with non-complex N-glycan extracellularly. As for hCG, no matter with or without the inhibition, shifts were only observed for all bands after the PNGase F digestion (white asterisks, Figure 3I) while all bands were Endo H resistant (yellow asterisks, Figure 3I). We also sought to investigated whether either KIF or SWA treatment affected the expression of non-glycosylated proteins. HLA-G can be only used in identifying the extravillous subtype of trophoblast because of its restricted expression [77] while cytokeratin 7 (KRT7) is used as a pan-trophoblast marker [78–80]; furthermore, zinc finger transcription factor GATA3 is a well-known regulator of trophoblast-specific gene expression and placental function, which is not expressed in syncytiotrophoblast [81–84]; therefore, GATA3 is used as a marker for mononuclear trophoblasts like EVT [78,85,86]. There is no present evidence that either KRT7 or GATA3 is N-glycosylated, whereas we found that their expression in JEG-3 cell were affected by the N-glycosylation as evidenced by lower levels of KRT7 and GATA3 in KIF-treated JEG-3 cell compared to those in control based on flow cytometric analyses of intact JEG-3 cell (Figures 3J-L). Collectively, these data demonstrate that the predominance of oligomannose type N-glycan in JEG-3 cell does not affect the secretion of glycoproteins but significantly reduces the expression of proteins related to trophoblast differentiation.

### 3.5 JEG-3 cells maintain core EVT-like properties despite pan-inhibition of sialylation

Although both KIF and SWA treatments induced similar changes in the binding of ConA, LEL, PHA-E, and PHA-L to JEG-3 cells, they exerted opposite effects on SNA and LCA binding. (Table 1). The vital role of sialylation in regulating the suppression of immune cell function has been demonstrated [7,8,60,87], we therefore questioned whether the reduced level of sialylation is the key in inducing phenotypic changes solely observed in KIF-treated JEG-3 cell. We used Neu5Ac analogue 3F_AX_-peracetyl-Neu5Ac (3FN), which has been demonstrated to inhibit the activity of sialyltransferases in a donor substrate competitive manner [88], to investigate the impact of reduced level of sialylation on JEG-3 cell phenotype and function. The 3FN treatment induced a significant reduction in α-2,6-sialylation (SNA) accompanied with increases of both β-1,6- and bisecting β-1,4-GlcNAc branches (PHA-L and -E, Figure 4A), suggesting an antagonistic relationship between terminal sialylation and GlcNAc branching, though it did not impair the proliferation of JEG-3 cell (Figure 4B). Levels of unfolded HLA-G fragment and hCG secretion (Figures 4C, D) were unaffected by the 3FN treatment in despite of the glycomic alteration. However, the GATA3 expression in JEG-3 cell was up-regulated by the 3FN treatment (Figure 4E). Combined with results from KIF and SWA treatment, we found that JEG-3 cell is losing its core EVT-like properties only when oligomannose type N-glycan predominates (Table 1).

**Figure 4.**
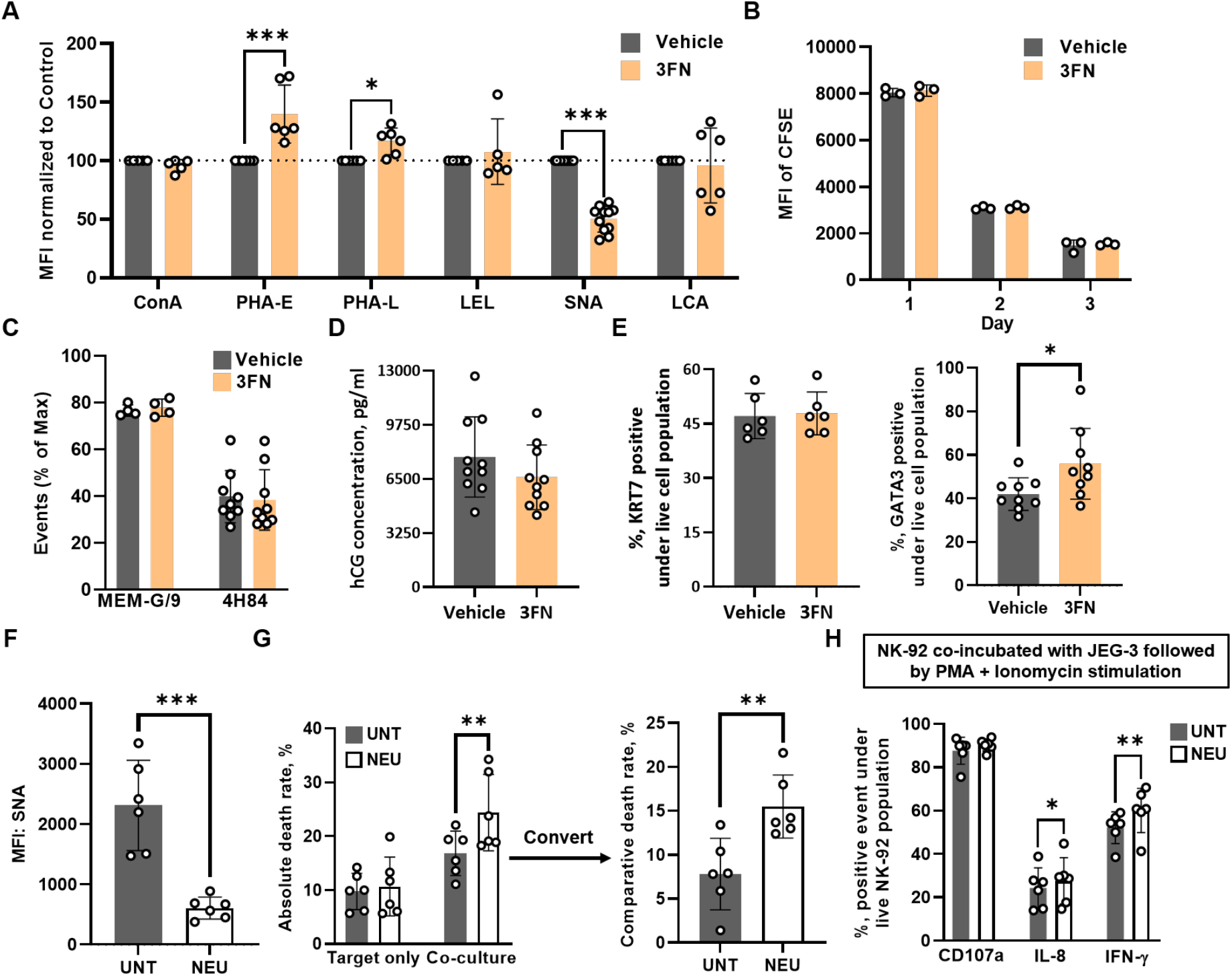
The role of sialylation in determining JEG-3 cell phenotype is not as decisive as its ability to generate hybrid type N-glycan. Phenotypes of JEG-3 cell with or without 3FN-induced inhibition of sialyltransferase activity were compared in terms of **(A)** flow cytometry of control and 3FN-treated JEG-3 cells using staining of representative lectins to detect the glycophenotype of the cell surface (N = 5-11), **(B)** cell proliferation over three-day incubation (N = 3), **(C)** levels of HLA-G1 on the cell surface or unfolded HLA-G fragment inside cell (N = 4 or 9), **(D)** whole hCG content in culture media analysed by ELISA (N =10), and **(E)** the proportion of KRT7 or GATA3 positive population in JEG-3 cells (N = 6 or 9). **(F)** Neuraminidase treatment of JEG-3 cell decreases the level of α-2,6-sialylation on its cell surface. N = 6. **(G)** The comparative death rates of untreated and neuraminidase-treated JEG-3 cell after co-culture with five-fold of NK-92 cell. N = 6. **(H)** Comparison in CD107a, IL-8, and IFN-γ expression by NK-92 cell under different preconditions. Four hours before the end of co-culture with either untreated or neuraminidase-treated JEG-3 cells, NK-92 cell were subjected to the stimulation of PMA/ionomycin. N = 6. Either paired Student’s t-test or Wilcoxon signed-rank test was used to compare the means between vehicle control and 3FN- or neuraminidase-treated JEG-3 cells. 3FN, 3F_AX_-Peracetyl Neu5Ac; NEU, neuraminidase; UNT, untreated.

**Table 1.** Glycomic and phenotypic alteration of JEG-3 cell after inhibitor treatment.

| Compared to control | Lectin binding to JEG-3 cell surface |  |  |  |  |  | Glycoproteins and differentiation markers |  |  |  |  |
| --- | --- | --- | --- | --- | --- | --- | --- | --- | --- | --- | --- |
|  | *ConA* | *PHA-E* | *PHA-L* | *LEL* | *SNA* | *LCA* | *HLA-G1* | *HLA-G fragment* | *hCG secretion* | *GATA3* | *KRT7* |
| **Inhibition of de-mannosylation** |  |  |  |  |  |  |  |  |  |  |  |
| **KIF** | ↑ | ↓ | ↓ | ↑ | ↓ | ↓ | N.S. | ↑ | ↑ | ↓ | ↓ |
| **SWA** |  |  |  |  | ↑ | ↑ | N.S. | N.S. | N.S. | N.S. | N.S. |
| **Inhibition of sialylation** |  |  |  |  |  |  |  |  |  |  |  |
| **3FN** | N.S. | ↑ | ↑ | N.S. | ↓ | N.S. | N.S. | N.S. | N.S. | ↑ | N.S. |

**Note:** Arrows denoted increased (↑), or decreased (↓) expression or abundance compared to the control. N = 6-12. N.S., non-significant.

### 3.6 Sialylation is important in protecting JEG-3 cell from NK-92 cell cytotoxicity

We further studied whether the de-sialylation could affect the resistance of JEG-3 cell against NK-92-mediated cytolysis (NK-resistance). Considering that the 3FN treatment cannot specifically reduce the level of sialylation (Figure 4A), we used neuraminidase to treat JEG-3 cells so that we could study the influence of sialylation level *per se* on the NK-resistance of JEG-3 cells. The neuraminidase treatment effectively reduced the level of α-2,6-sialylation (SNA) on JEG-3 cell surface (Figure 4F) without impairing the vitality of JEG-3 cell (Figure 4G). JEG-3 cell with or without neuraminidase treatment was subjected to cytotoxicity assays (Supplementary Figure 3A), and de-sialylated JEG-3 cell exhibited an impaired NK-resistance indicated by an increased comparative cell death after the co-incubation with NK-92 cells (Figure 4G and Supplementary Figure 3B). We further studied whether NK-92 cells become more cytotoxic when co-cultured with de-sialylated JEG-3 cells, and we determined the NK-92 cell cytotoxicity after co-culture with JEG-3 cell by measuring the expression of NK cell degranulation marker CD107a [89], and the generation of cytokines IL-8 and IFN-γ which have been reported to exert inhibitory effects on trophoblast invasion [90–92]. Minimal expression of CD107a and production of IL-8 and IFN-γ by NK-92 cell was observed after 10-hour co-culture; however, NK-92 cell exhibited a high expression of CD107a and generation of IL-8 and IFN-γ upon stimulation of PMA and ionomycin (Supplementary Figure 4). Statistical analysis revealed that, NK-92 cells that were stimulated with PMA/Ionomycin after first being co-cultured with JEG-3 cells showed a significantly reduced IL-8 and IFN-g response compared to NK-92 cells similarly stimulated following co-culture with neuraminidase treated JEG-3 cells, indicating that sialylation impacts the JEG-3 immunomodulation of NK-92 cells (Figure 4H).

### 3.7 The immunomodulatory effect of JEG-3 cell secretome on NK-92 cell is only affected when oligomannose type N-glycans predominate

Membrane glycoproteins expressed by EVT have been reported to interact with receptors derived from decidual immune cells, educating the latter to an immunosuppressive phenotype [37–39,93]. Likewise, secretory glycoproteins of EVT have also been reported to promote the peripheral immune cell differentiation towards an immunosuppressive phenotype [46,94,95]. We therefore sought to investigate whether and to what extent the glycan remodelling affects immunomodulation of JEG-3 cell on immune cell with cytotoxicity, and we evaluated this effect from two aspects: the cell-cell interaction and secretome (Supplementary Figure 3A). We first observed that NK-92 cells exhibited comparable cytotoxicity towards K562 cell when pre-conditioned by either co-culture with JEG-3 cells or by matched supernatant (Figure 5A). To study whether the immunomodulation of JEG-3 cells on NK-92 cell cytotoxicity through cell-cell interaction is affected by the glycan remodelling, NK-92 cells were pre-conditioned by co-culture with JEG-3 cells possessing different glycomic profiles before co-culture with K562 cells (Supplementary Figure 3A). It showed that the interaction with any inhibitor-treated JEG-3 cell did not affect the NK-92 cell cytotoxicity towards K562 cell when compared to control (Figure 5B). As for verifying the immunomodulation of JEG-3 cells NK-92 cell cytotoxicity through secretory glycoproteins, NK-92 cells were pre-conditioned with supernatant from different inhibitor-treated JEG-3 cells. Repressed cytotoxicity of NK-92 cell towards K562 cell was only observed in NK-92 pre-conditioned with supernatant of KIF-treated JEG-3 cell (SN-KIF, Figure 5C). NK-92 cell pre-conditioned with different calibrated culture media indicated that the remaining drugs in culture media has limited effect on the NK-92 cell cytotoxicity, except for an enhanced cytotoxicity observed in NK-92 cells pre-conditioned with culture media supplemented with 3FN (CM-3FN, Figure 5D). We further found that NK-92 cells pre-conditioned with supernatant of JEG-3 cells exhibited an enhanced cytotoxicity to K562 cell, yet this enhancement can be reversed by sialic acid (Figure 5E), highlighting that secretory glycoproteins derived from JEG-3 cells play a major role in affecting NK-92 cell cytotoxicity.

**Figure 5.**
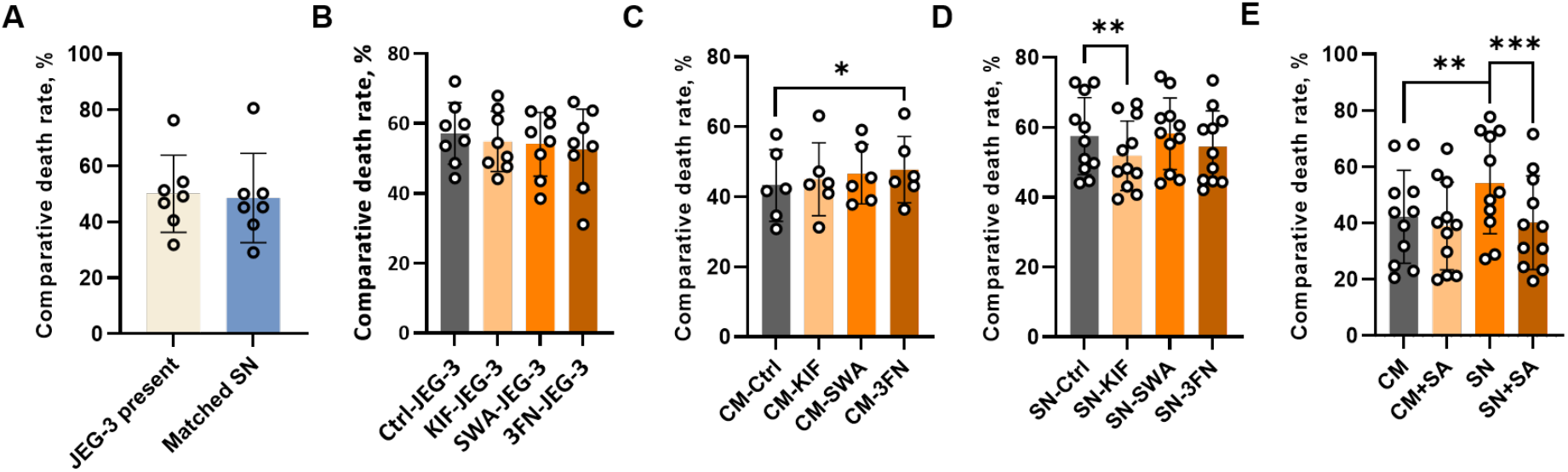
The immunomodulatory effect of secreted glycoproteins derived from JEG-3 cells can be modified through glycomic remodelling. The comparative death rates of K562 cells after 10 h co-culture with equal number of NK-92 cells pre-conditioned with **(A)** JEG-3 cell monolayer or matched SN, **(B)** exposure to JEG-3 cells with or without different types of glycan remodelling, **(C)** CM in the absence or presence of different inhibitors, **(D)** SN of JEG-3 cells with or without different types of glycan remodelling, or **(E)** CM or SN with or without SA supplement. Either paired Student’s t-test or Wilcoxon signed-rank test was used to compare the means between paired groups. N = 6-12. CM, culture media; SN, supernatant; SA, sialic acid.

## 4 Discussion

The goal of this study was to verify the regulatory role of N-glycosylation in shaping immunological properties of choriocarcinoma cell line JEG-3 which was used as an *in vitro* model of EVT. Pharmacological inhibition of α-mannosidase II caused complete loss of GlcNAc branching in JEG-3 cell but triggered compensatory increases in poly-LacNAc, α-2,6-sialylation, and core α-1,6-fucosylation, without affecting EVT marker expression, hCG secretion, or its immunomodulation on NK-92 cell. In contrast, inhibition of α-mannosidase I led to the predominance of immature oligomannose type N-glycans and resulted in marked phenotypic alterations, including intracellular accumulation of unfolded HLA-G fragment, reduced expression of GATA3 and KRT7, increased hCG secretion, and inadequate immunomodulation on NK-92 cells. Our model indicates that alterations in the EVT glycomic profile may lead to altered immunomodulation at the maternal-fetal interface, providing insight into why certain N-glycomic profiles are linked to pathological placentae [21–28], and offering targets for drug therapy in pregnancy.

Our findings that the SWA treatment induced additional poly-LacNAc extension in choriocarcinoma cell lines JEG-3 and JAR but not in erythroleukemia cell line K562 (Figures 1B, 2A, and 2B) implied that the capability of generating bioequivalent glycans when α-mannosidase II is inhibited might be an intrinsic property for specific cell type. Similar compensatory responses to N-glycosylation inhibition have also been reported in transgenic mice [96,97]. For example, Mkhikian *et al* found that immune cells in T-cell-specific-*Mgat2*-deficient mice lacking N-glycan branching enzymes present glycans on their surface formed of LacNAc arranged in chains, which are biologically equivalent to LacNAc arranged in branches, allowing the immune cells to behave as normal [96]. They further demonstrated a self-correcting mechanism that those enzymes catalysing GlcNAc branching are located earlier in the Golgi apparatus than those catalysing the generation of poly-LacNAc extension; therefore, if the branching enzymes fail to add LacNAc subunits to potential GlcNAc branches, the extending enzymes can step in later to add the missing components [96]. JEG-3 and JAR cells likely employ this mechanism, compensating for the absence of GlcNAc branching on the α-1,6-Man arm with enhanced poly-LacNAc extension on the α-1,3-Man arm (Figure 1A), thereby maintaining LacNAc homeostasis. Interestingly, JEG-3 cells displayed a more pronounced increase in poly-LacNAc extension following SWA treatment compared with JAR cells (Figure 2C). Together with previous glycomic analyses showing that primary EVT cells possess higher levels of poly-LacNAc extension than villous trophoblasts under physiological conditions [64], these findings suggest that poly-LacNAc structures may play an important role in the regulation of EVT function. More importantly, this glycan feature may provide a potential biomarker for distinguishing pathological EVT from healthy EVT and for identifying disease-associated N-glycan structures.

Specific glycan structures have been linked to EVT phenotypes, with bisecting β-1,4-GlcNAc promoting invasion and β-1,6-GlcNAc branching inhibiting it [29,31,98]. A previous study found an increase in the oligomannose type N-glycans in chorionic villi from women with early-onset preeclampsia and a decreased hCG production by chorionic villi by measuring supernatants of histoculture [56]. However, our results indicate that EVT phenotype of JEG-3 cells is not solely determined by glycan abundance. SWA- or 3FN-induced glycan remodelling did not affect HLA-G1 expression or hCG secretion (Figures 3C, 3H, 4C, and 4D), and HLA-G1 remained expressed even when decorated predominantly with oligomannose N-glycans after KIF treatment (Figures 3C-F). This suggests functional redundancy in N-glycosylation, whereby glycoproteins bearing improper N-glycans can still be correctly folded and expressed on cell surface or secreted. Although aberrant hCG glycosylation has been linked to adverse pregnancy outcomes [25,27,99], the hCG secretion of SWA-treated JEG-3 cell was unaffected and there was even an increased secretion by KIF-treated samples (Figure 3H), suggesting that the glycosylation of hCG is not directly related to its production. In fact, free hCGα purified from the culture medium of SWA-treated JEG-3 cells has been reported to combine with hCGβ with higher avidity [100]. It is possible that subunits hCGα derived from KIF-treated JEG-3 cells are abundantly decorated with oligomannose N-glycans, of which conformation and size is optimized for its combination with hCGβ, eventually results in higher production and the concomitant secretion. Notably, NK-92 cells pre-conditioned with SN-KIF showed suppressed killing on K562 cells (Figure 5D). The increased hCG secretion by KIF-treated JEG-3 cell (Figure 3H) might contribute to the suppressed cytotoxicity of NK-92 cell, but this suppression also suggested that the simplification of N-glycan heterogeneity might be related to the immunosuppressive properties of trophoblast-derived glycoproteins. GATA3 is a protein without N-glycosylation site (ref. GlyGen P23771-1) and its expression has been related to the maturation of placenta as indicated by a higher expression of GATA3 observed in immature placentae when compared to that in mature placentae [81]. We observed that GATA3 expression in JEG-3 cell is sensitive to both inhibition of α-mannosidase I (Figures 3L) and pan-inhibition of sialyltransferases (Figures 4E), indicating that altered N-glycosylation occurring during pregnancy can affect GATA3 expression in trophoblast cells and likely impact their development. Cao et al have demonstrated that N-glycosylation pathway activity is significantly enhanced during the differentiation of human trophoblast stem cells into EVTs [101]. Future studies are warranted to define the role of N-glycosylation in this differentiation process, thereby improving our understanding of how EVTs acquire their unique immunomodulatory functions.

We also observed that in response to SWA treatment JEG-3 cells exhibited a higher level of α-2,6-sialosides than the control (Figure 1B), which might be fulfilled through capping β-1,2- or β-1,4-GlcNAc branches attached to α-1,3-Man arm with sialic acids, while JAR cells retain a comparable level of α-2,6-sialosides on the cell surface as the control (Figure 2A). Sialylated antigen glycans derived from trophoblasts have been reported to be involved in the mediation of B cell suppression to establish feto-maternal tolerance in mice [57]. Another deficient mouse model also highlighted the pivotal role of sialic acid in fetal-maternal immune homeostasis during pregnancy [58]. EVTs from patients with recurrent pregnancy loss has been reported to exhibit reduced sialylation, coinciding with an increased proportion of Siglec-7 positive decidual dNK cells, implying that the disruption of a key immunological checkpoint that normally promotes EVT invasion induced by defective EVT sialylation potentially contributes to recurrent pregnancy loss [102]. In this study, we have investigated the immunomodulatory effect of secreted and membrane glycoproteins from JEG-3 cells with glycan remodelling, modelling EVT-mediated immunomodulation via soluble immunomodulators and cell-cell interactions. Although neuraminidase treatment rendered JEG-3 cells more susceptible to NK-cell-mediated responses (Figure 4G), neither exposure of NK-92 cells to SN-3FN nor co-culture with 3FN-treated JEG-3 cells affected their cytolytic activity against K562 cells. Likewise, SWA-treated JEG-3 cells, which displayed significantly increased surface sialylation (Figure 1B), failed to modulate NK-92 cytotoxicity (Figure 5B, D). These findings suggest that alterations in JEG-3 cell sialylation alone are insufficient to account for the observed changes in immunomodulation of NK-92 cell. A limitation of our study was that decidual NK cell from pregnant women without complications was not available for cytotoxicity assays. It is therefore worth investigating whether pathological EVTs that fail to silence the cytotoxic potential of decidual NK cells are possessing a decreased level of sialylation. We also found that, instead of the cytotoxicity being suppressed, NK-92 cell exhibited enhanced killing on K562 cell when pre-conditioned with supernatant of JEG-3 cell compared to those with matched calibrated culture media, and this enhancement can be reversed by the addition of sialic acid (Figure 5E), again highlighting the immunomodulatory effect of sialic acid on NK-92 cell cytotoxicity [103]. Together, these findings underscore the importance of sialoside in mediating NK cell cytotoxicity, but also question the manner that JEG-3 cell adopts to exert immunomodulatory effect through cell surface expression of sialylated antigens or secretion of sialylated glycoprotein.

Taken together, our findings suggest that the expression of N-glycan components poly-LacNAc extension and sialoside is required for JEG-3 cell to perform immunomodulation on NK-92 cells and its immunological properties are affected only under predominance of oligomannose type N-glycan. The insight gained from our *in vitro* model will aid the direction of future research on how N-glycosylation affects primary EVTs exerting immunomodulatory effect on decidual immune cells during early pregnancy. Further larger cohorts should be recruited for studies on phenotype and function of glycoproteins derived from the interstitial population of EVT, with a focus on their ligation with receptors expressed on decidual NK cell. Furthermore, glycosylation-induced modulation of EVT immunological properties is pathologically relevant, such as in the development of neoplastic glycosylation patterns that support cancer metastasis, and research into how to modulate EVTs N-glycosylation during pregnancy will support future clinical advances.

## Author Contributions

This research was designed by ZH and MRJ. All experiments were performed by ZH. The obtained data were analysed by ZH, ATHC, and XF. These results were interpreted by ZH, ATHC, GW, XF, and MRJ. The manuscript was written by ZH and ATHC. MRJ contributed to all subsequent editing and approved the final submitted draft.

---

**Supplementary Figure 1.**
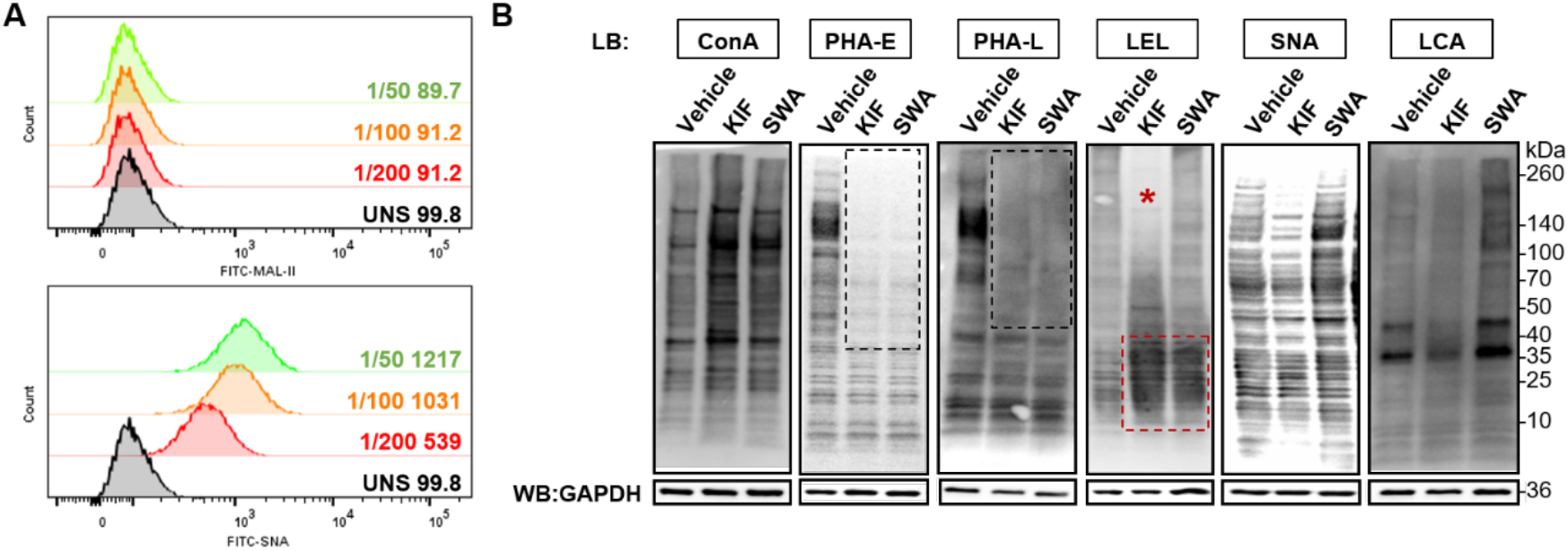
**(A)** Representative histograms of MAL II or SNA staining of wildtype JEG-3 cells under different doses of lectin added. UNS, unstained control. Representative histograms are shown. **(B)** Lectin blotting on whole cell lysates of both inhibitor-treated JEG-3 cells and vehicle control. Decreased bindings of both PHA-E and -L were restricted to glycoproteins with molecular weight of more than 40 kDa (black dotted box). A global increase of ConA binding was observed in both inhibitor-treated populations; meanwhile, an increased LEL binding in glycoproteins below 40 kDa was observed in both populations (red dotted box); however, glycoproteins of which molecular weight higher than ∼70 kDa were devoid of LEL binding in KIF-treated JEG-3 cells only (red asterisk). Representative blots are shown.

**Supplementary Figure 2.**
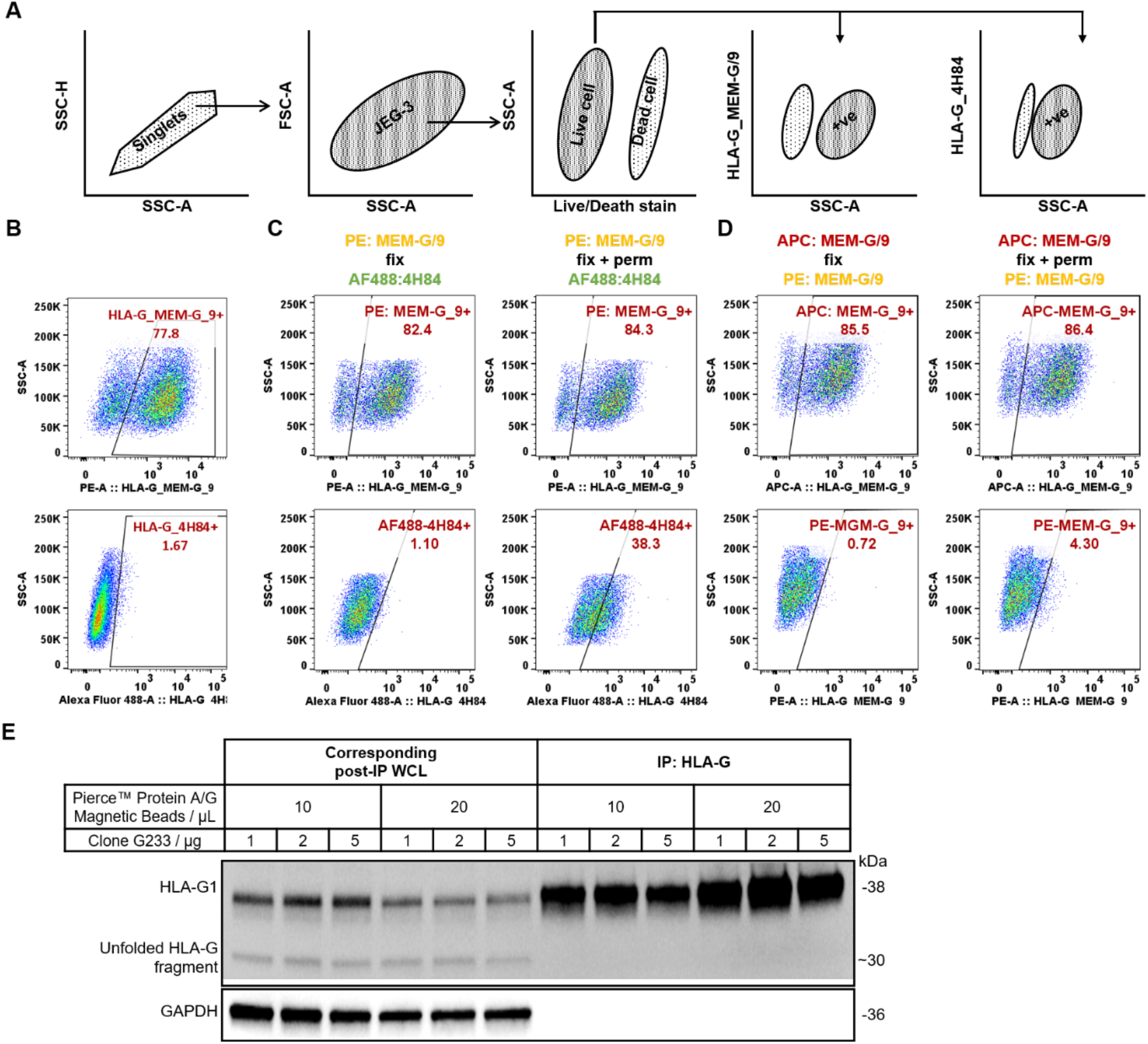
**(A)** The gating strategy applied for sorting out desired JEG-3 cell population. SSC-A against SSC-H is first used to gate in all singlets, and then SSC-A against FSC-A plot is further used to determine the size and complexity of desired JEG-3 cells population. After the exclusion of dead cells, either MEM-G/9 or 4H84 positive population was plotted under live cell population. **(B)** Representative scatter plot of JEG-3 cell surface staining with different clones of anti-HLA-G mAb. Intact JEG-3 cells were stained with either PE-MEM-G/9 or AF488-4H84 in 1:100 antibody dilution (1 µl of antibody stock in 100 µl of cell suspension, same as below unless otherwise indicated). **(C)** JEG-3 cell can only be stained by clone 4H84 after permeabilization. Intact JEG-3 cells were first stained with PE-MEM-G/9, subjected to fixation and/or permeabilization, and eventually incubated with AF488-4H84. **(D)** Clone MEM-G/9 can only bind to HLA-G expressed on cell surface. JEG-3 cells were first saturated with 10 µl of antibody stock in 100 µl of cell suspension of APC-MEM-G/9, subjected to fixation and/or permeabilization, and eventually incubated with PE-MEM-G/9. Those gates for determining MEM-G/9 positive population among groups were identical, and so were those gates for determining 4H84 positive population. Representative scatter plots are shown. **(E)** The immunoprecipitation of HLA-G1 proteins from wildtype JEG-3 whole cell lysate followed by Western blotting of all HLA-G isoforms. Different combinations of Pierce™ Protein A/G Magnetic Beads (10 and 20 µl) and clone G233 (1, 2 and 5 µg) were used to immunoprecipitate native HLA-G1 protein. Afterwards, both immunoprecipitated HLA-G (IP-HLA-G) and whole cell lysate after immunoprecipitation (post-IP WCL) were analysed by Western blotting using clone 4H84. The Western blot of GAPDH was used to indicate the specificity of clone G233 in pulling down HLA-G1 molecules.

**Supplementary Figure 3.**
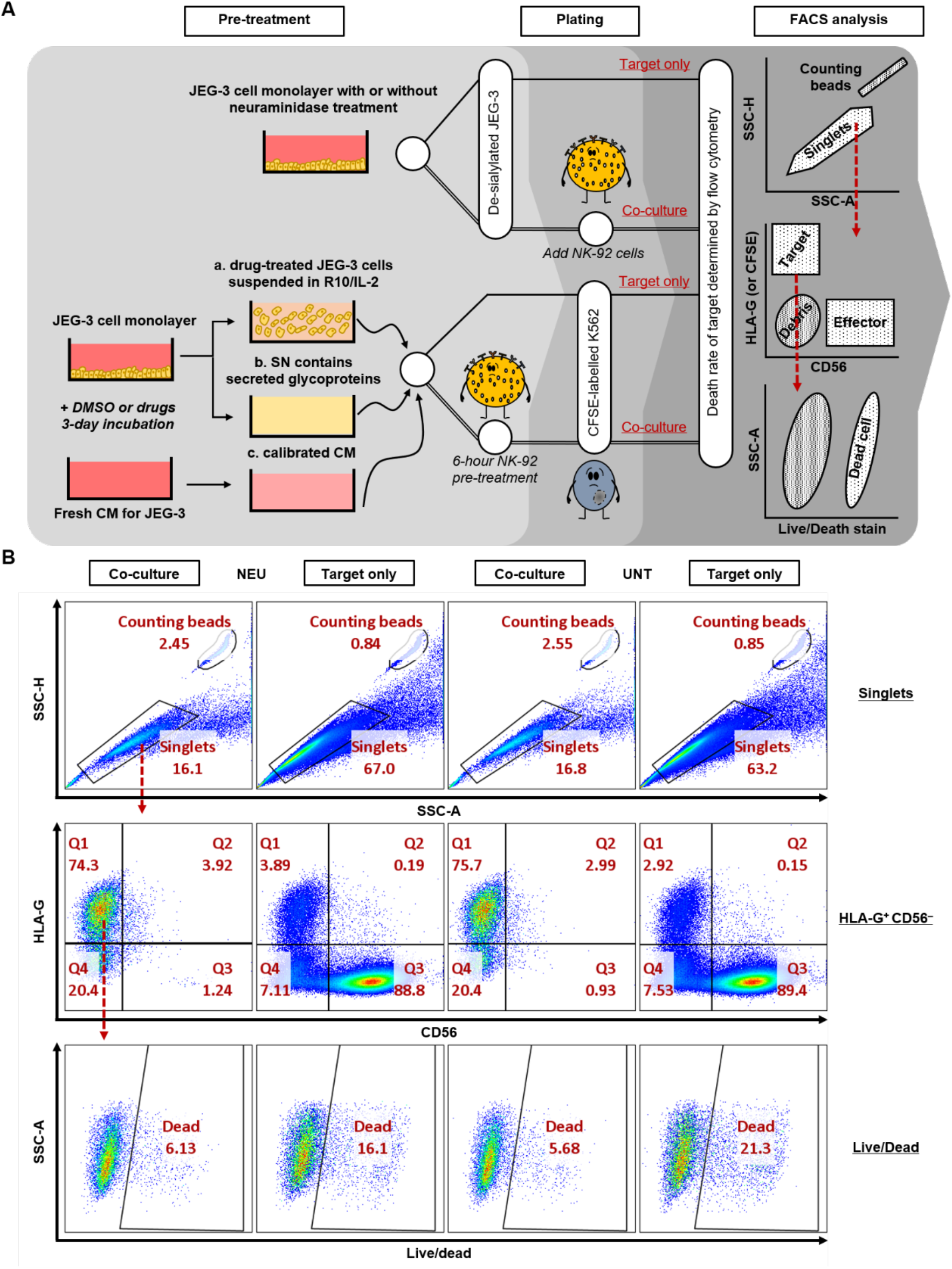
**(A)** The workflow of flow cytometry-based cytotoxicity assay and gating strategy to determine the absolute death rate of target in flow cytometric analyses, in which either JEG-3 or K562 cells are the target while NK-92 cells are the effector. A consistent number of JEG-3 cells were seeded in two separate wells, allowing them to recover for at least 8 hours until adhesion reforms before co-incubating with NK-92 cells. RPMI 1640 medium complemented with 10% FBS and 200 IU/ml IL-2 (R-10 + IL-2) was used as culture media for co-culture of JEG-3 and NK-92 cells. JEG-3 cell monolayer was first treated with neuraminidase and then five-fold of NK-92 cell is added to the “Co-culture” well so that the E:T ratio would be approximately 5:1, while the “Target only” well seeded with de-sialylated JEG-3 cells was setup to determine their spontaneous death. Same setting was used for the co-culture of K562 and NK-92 cells except that the effector was pre-conditioned with either different JEG-3 cell suspension, different used culture media of JEG-3 cells, or calibrated culture media for JEG-3 cells supplemented with different inhibitors. K562 was labelled with CFSE before the cytotoxicity assay. Doublets and debris were excluded and then markers HLA-G (or CFSE) and CD56 were used to discriminate targets from effectors. Target populations were gated and then live cells were discriminated against dead cells. The percentage of live/dead stain positive population under target population indicated the absolute death rate of either “Target only” (spontaneous death) or “Co-culture” (experimental death) arms. **(B)** Gating strategy that is used to determine the absolute death of target cells. Doublets were first excluded in SSC-H-SSC-A plot, and then the dead target (live/dead stain positive) population was plotted out under Q1 quadrant (HLA-G^+^ CD56^-^ population). The proportion of counting bead was plotted in in SSC-H-SSC-A plot to indicate cell numbers used in each group.

**Supplementary Figure 4.**
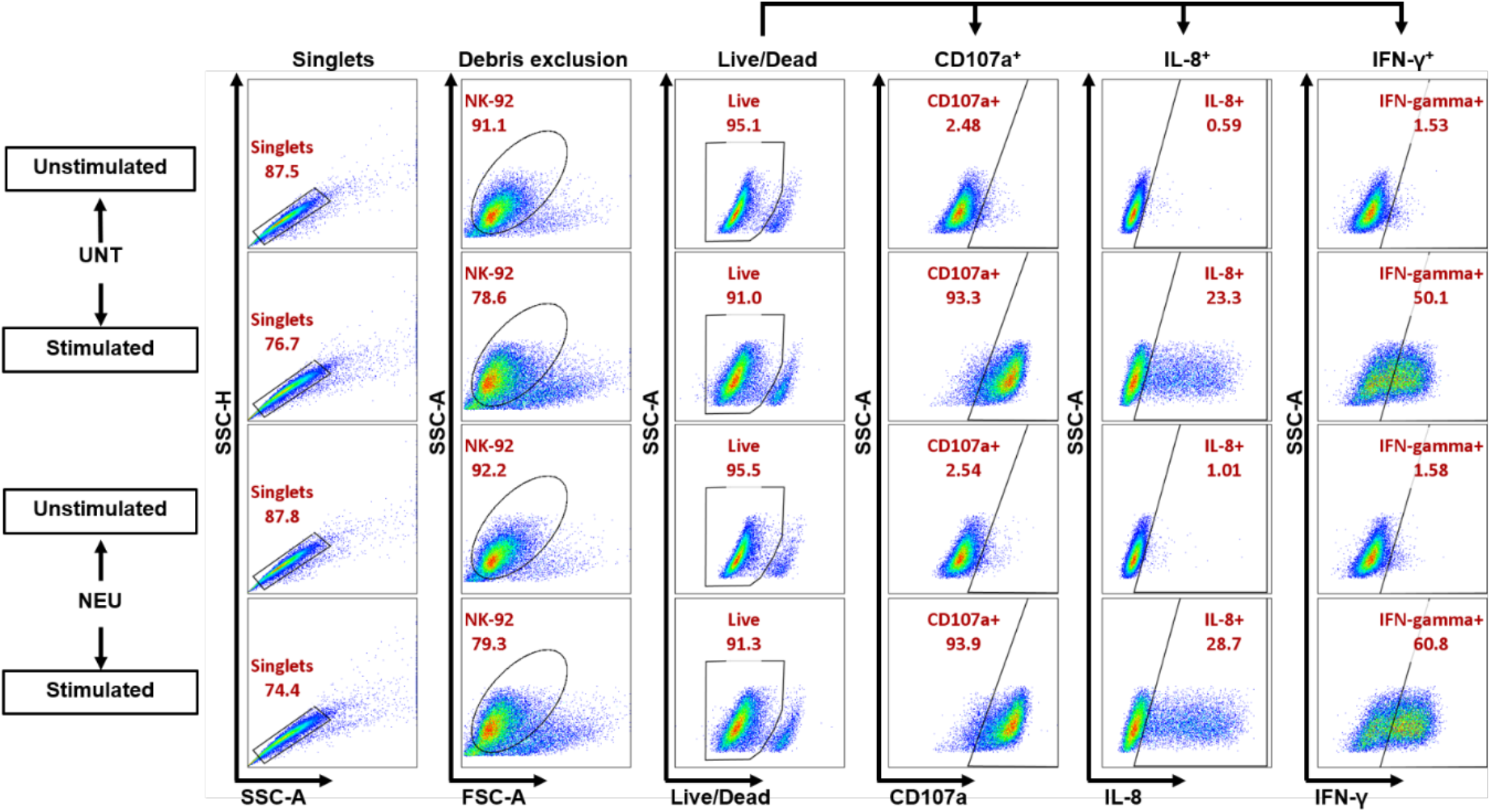
Gating strategy used to determine the level of CD107a, IL-8 and IFN-γ under live NK-92 population. Doublets were first excluded under SSC-A-SSC-H plot and then debris were plotted out under singlets in FSC-A-SSC-A plot. Dead populations were excluded so that the levels of CD107a, IL-8 and IFN-γ positive population were determined under live NK-92 singlets.

## References

1. Fisher, P.; Thomas-Oates, J.; Wood, A.J.; Ungar, D. The N-Glycosylation Processing Potential of the Mammalian Golgi Apparatus. Front Cell Dev Biol 2019, 7, 157, doi:10.3389/fcell.2019.00157.

2. Hirata, T.; Kizuka, Y. N-Glycosylation. In The Role of Glycosylation in Health and Disease, Lauc, G., Trbojević-Akmačić, I., Eds.; Springer International Publishing: Cham, 2021; pp. 3–24.

3. Aebi, M. N-linked protein glycosylation in the ER. Biochim Biophys Acta 2013, 1833, 2430–2437, doi:10.1016/j.bbamcr.2013.04.001.

4. Hao, C.; Zou, Q.; Bai, X.; Shi, W. Effect of glycosylation on protein folding: From biological roles to chemical protein synthesis. iScience 2025, 28, 112605, doi:10.1016/j.isci.2025.112605.

5. Almahayni, K.; Spiekermann, M.; Fiore, A.; Yu, G.; Pedram, K.; Möckl, L. Small molecule inhibitors of mammalian glycosylation. Matrix Biol Plus 2022, 16, 100108, doi:10.1016/j.mbplus.2022.100108.

6. Song, X.; Zhou, Z.; Li, H.; Xue, Y.; Lu, X.; Bahar, I.; Kepp, O.; Hung, M.C.; Kroemer, G.; Wan, Y. Pharmacologic Suppression of B7-H4 Glycosylation Restores Antitumor Immunity in Immune-Cold Breast Cancers. Cancer Discov 2020, 10, 1872–1893, doi:10.1158/2159-8290.Cd-20-0402.

7. Yoshimura, M.; Ihara, Y.; Ohnishi, A.; Ijuhin, N.; Nishiura, T.; Kanakura, Y.; Matsuzawa, Y.; Taniguchi, N. Bisecting N-acetylglucosamine on K562 cells suppresses natural killer cytotoxicity and promotes spleen colonization. Cancer research 1996, 56, 412–418.

8. Hudak, J.E.; Canham, S.M.; Bertozzi, C.R. Glycocalyx engineering reveals a Siglec-based mechanism for NK cell immunoevasion. Nature Chemical Biology 2014, 10, 69–75, doi:10.1038/nchembio.1388.

9. Zhuang, X.; Woods, J.; Ji, Y.; Scheich, S.; Mo, F.; Rajagopalan, S.; Coulibaly, Z.A.; Voss, M.; Urlaub, H.; Staudt, L.M.;, et al. Functional genomics identifies N-acetyllactosamine extension of complex N-glycans as a mechanism to evade lysis by natural killer cells. Cell Rep 2024, 43, 114105, doi:10.1016/j.celrep.2024.114105.

10. Becker, J.C.; Andersen, M.H.; Schrama, D.; Thor Straten, P. Immune-suppressive properties of the tumor microenvironment. Cancer Immunology, Immunotherapy 2013, 62, 1137–1148, doi:10.1007/s00262-013-1434-6.

11. Stowell, S.R.; Ju, T.; Cummings, R.D. Protein Glycosylation in Cancer. Annual Review of Pathology: Mechanisms of Disease 2015, 10, 473–510, doi:10.1146/annurev-pathol-012414-040438.

12. Fry, S.A.; Afrough, B.; Lomax-Browne, H.J.; Timms, J.F.; Velentzis, L.S.; Leathem, A.J. Lectin microarray profiling of metastatic breast cancers. Glycobiology 2011, 21, 1060–1070, doi:10.1093/glycob/cwr045.

13. Häuselmann, I.; Borsig, L. Altered tumor-cell glycosylation promotes metastasis. Frontiers in oncology 2014, 4, 28.

14. Jandus, C.; Boligan, K.F.; Chijioke, O.; Liu, H.; Dahlhaus, M.; Démoulins, T.; Schneider, C.; Wehrli, M.; Hunger, R.E.; Baerlocher, G.M. Interactions between Siglec-7/9 receptors and ligands influence NK cell–dependent tumor immunosurveillance. The Journal of Clinical Investigation 2014, 124, 1810–1820.

15. Tilburgs, T.; Crespo Â, C.; van der Zwan, A.; Rybalov, B.; Raj, T.; Stranger, B.; Gardner, L.; Moffett, A.; Strominger, J.L. Human HLA-G+ extravillous trophoblasts: Immune-activating cells that interact with decidual leukocytes. Proceedings of the National Academy of Sciences 2015, 112, 7219–7224, doi:10.1073/pnas.1507977112.

16. Turco, M.Y.; Moffett, A. Development of the human placenta. Development 2019, 146, dev163428.

17. Wang, X.Q.; Li, D.J. The mechanisms by which trophoblast-derived molecules induce maternal-fetal immune tolerance. Cell Mol Immunol 2020, 17, 1204–1207, doi:10.1038/s41423-020-0460-5.

18. Knöfler, M.; Haider, S.; Saleh, L.; Pollheimer, J.; Gamage, T.; James, J. Human placenta and trophoblast development: key molecular mechanisms and model systems. Cell Mol Life Sci 2019, 76, 3479–3496, doi:10.1007/s00018-019-03104-6.

19. Rizzuto, G.; Erlebacher, A. Trophoblast antigens, fetal blood cell antigens, and the paradox of fetomaternal tolerance. J Exp Med 2022, 219, doi:10.1084/jem.20211515.

20. Wang, T.; Wang, Y.; Wu, Y.; Zhang, Y.; Ma, F. The dynamics of glycosylation in trophoblast cells: development, function, and pregnancy-related disorders. Ann Med 2026, 58, 2593420, doi:10.1080/07853890.2025.2593420.

21. Marini, M.; Bonaccini, L.; Thyrion, G.D.; Vichi, D.; Parretti, E.; Sgambati, E. Distribution of sugar residues in human placentas from pregnancies complicated by hypertensive disorders. Acta Histochem 2011, 113, 815–825, doi:10.1016/j.acthis.2010.12.001.

22. Sukhikh, G.T.; Ziganshina, M.M.; Nizyaeva, N.V.; Kulikova, G.V.; Volkova, J.S.; Yarotskaya, E.L.; Kan, N.E.; Shchyogolev, A.I.; Tyutyunnik, V.L. Differences of glycocalyx composition in the structural elements of placenta in preeclampsia. Placenta 2016, 43, 69–76, doi:10.1016/j.placenta.2016.05.002.

23. Tannetta, D.; Masliukaite, I.; Vatish, M.; Redman, C.; Sargent, I. Update of syncytiotrophoblast derived extracellular vesicles in normal pregnancy and preeclampsia. J. Reprod. Immunol. 2017, 119, 98–106, doi:10.1016/j.jri.2016.08.008.

24. Kobayashi, Y.; Masuda, K.; Banno, K.; Kobayashi, N.; Umene, K.; Nogami, Y.; Tsuji, K.; Ueki, A.; Nomura, H.; Sato, K. Glycan profiling of gestational choriocarcinoma using a lectin microarray. Oncology reports 2014, 31, 1121–1126, doi:10.3892/or.2014.2979.

25. Elliott, M.M.; Kardana, A.; Lustbader, J.W.; Cole, L.A. Carbohydrate and peptide structure of the α- and β-subunits of human chorionic gonadotropin from normal and aberrant pregnancy and choriocarcinoma. Endocrine 1997, 7, 15–32, doi:10.1007/BF02778058.

26. Zhang, M.; Wang, M.; Gao, R.; Liu, X.; Chen, X.; Geng, Y.; Ding, Y.; Wang, Y.; He, J. Altered β1,6-GlcNAc and bisecting GlcNAc-branched N-glycan on integrin β1 are associated with early spontaneous miscarriage in humans. Human Reproduction 2015, 30, 2064–2075, doi:10.1093/humrep/dev153.

27. Ibeto, L.; Antonopoulos, A.; Grassi, P.; Pang, P.C.; Panico, M.; Bobdiwala, S.; Al-Memar, M.; Davis, P.; Davis, M.; Norman Taylor, J.;, et al. Insights into the hyperglycosylation of human chorionic gonadotropin revealed by glycomics analysis. PLoS One 2020, 15, e0228507, doi:10.1371/journal.pone.0228507.

28. Wang, H.; Fan, N.; Cui, X.; Xie, R.; Tang, Y.; Thomas, A.M.; Li, S.; Zhang, J.V.; Liu, S.; Qin, H. BMP5 promotes trophoblast functions upon N-glycosylation via the BMP5-SMAD1/5 signaling pathway in preeclampsia. Placenta 2024, doi:10.1016/j.placenta.2024.11.002.

29. Deng, Q.; Chen, Y.; Yin, N.; Shan, N.; Luo, X.; Tong, C.; Zhang, H.; Baker, P.N.; Liu, X.; Qi, H. N-acetylglucosaminyltransferase V inhibits the invasion of trophoblast cells by attenuating MMP2/9 activity in early human pregnancy. Placenta 2015, 36, 1291–1299, doi:10.1016/j.placenta.2015.08.014.

30. Niimi, K.; Yamamoto, E.; Fujiwara, S.; Shinjo, K.; Kotani, T.; Umezu, T.; Kajiyama, H.; Shibata, K.; Ino, K.; Kikkawa, F. High expression of N-acetylglucosaminyltransferase IVa promotes invasion of choriocarcinoma. Br J Cancer 2012, 107, 1969–1977, doi:10.1038/bjc.2012.496.

31. Yamamoto, E.; Ino, K.; Miyoshi, E.; Inamori, K.; Abe, A.; Sumigama, S.; Iwase, A.; Kajiyama, H.; Shibata, K.; Nawa, A.;, et al. N-acetylglucosaminyltransferase V regulates extravillous trophoblast invasion through glycosylation of alpha5beta1 integrin. Endocrinology 2009, 150, 990–999, doi:10.1210/en.2008-1005.

32. Liao, W.C.; Liu, C.H.; Chen, C.H.; Hsu, W.M.; Liao, Y.Y.; Chang, H.M.; Lan, C.T.; Huang, M.C.; Shyu, M.K. beta-1,4-Galactosyltransferase III suppresses extravillous trophoblast invasion through modifying beta1-integrin glycosylation. Placenta 2015, 36, 357–364, doi:10.1016/j.placenta.2015.01.008.

33. Deng, Q.; Liu, X.; Yang, Z.; Xie, L. Expression of N-Acetylglucosaminyltransferase III Promotes Trophoblast Invasion and Migration in Early Human Placenta. Reprod Sci 2019, 26, 1373–1381, doi:10.1177/1933719118765967.

34. Yu, M.; Cui, X.; Wang, H.; Liu, J.; Qin, H.; Liu, S.; Yan, Q. FUT8 drives the proliferation and invasion of trophoblastic cells via IGF-1/IGF-1R signaling pathway. Placenta 2019, 75, 45–53, doi:10.1016/j.placenta.2018.11.005.

35. Hoek, M.; Demmers, L.C.; Wu, W.; Heck, A.J.R. Allotype-Specific Glycosylation and Cellular Localization of Human Leukocyte Antigen Class I Proteins. Journal of Proteome Research 2021, 20, 4518–4528, doi:10.1021/acs.jproteome.1c00466.

36. McMaster, M.; Zhou, Y.; Shorter, S.; Kapasi, K.; Geraghty, D.; Lim, K.-H.; Fisher, S. HLA-G isoforms produced by placental cytotrophoblasts and found in amniotic fluid are due to unusual glycosylation. The Journal of Immunology 1998, 160, 5922–5928.

37. Tilburgs, T.; Evans, J.H.; Crespo Â C.; Strominger, J.L. The HLA-G cycle provides for both NK tolerance and immunity at the maternal-fetal interface. Proceedings of the National Academy of Sciences 2015, 112, 13312–13317, doi:10.1073/pnas.1517724112.

38. Fu, B.; Zhou, Y.; Ni, X.; Tong, X.; Xu, X.; Dong, Z.; Sun, R.; Tian, Z.; Wei, H. Natural killer cells promote fetal development through the secretion of growth-promoting factors. Immunity 2017, 47, 1100–1113. e1106.

39. Petroff, M.G.; Sedlmayr, P.; Azzola, D.; Hunt, J.S. Decidual macrophages are potentially susceptible to inhibition by class Ia and class Ib HLA molecules. Journal of Reproductive Immunology 2002, 56, 3–17.

40. Salvany-Celades, M.; van der Zwan, A.; Benner, M.; Setrajcic-Dragos, V.; Bougleux Gomes, H.A.; Iyer, V.; Norwitz, E.R.; Strominger, J.L.; Tilburgs, T. Three Types of Functional Regulatory T Cells Control T Cell Responses at the Human Maternal-Fetal Interface. Cell Reports 2019, 27, 2537–2547.e2535, 10.1016/j.celrep.2019.04.109.

41. Lee, C.L.; Guo, Y.; So, K.H.; Vijayan, M.; Guo, Y.; Wong, V.H.; Yao, Y.; Lee, K.F.; Chiu, P.C.; Yeung, W.S. Soluble human leukocyte antigen G5 polarizes differentiation of macrophages toward a decidual macrophage-like phenotype. Human Reproduction 2015, 30, 2263–2274, doi:10.1093/humrep/dev196.

42. Martayan, A.; Sibilio, L.; Setini, A.; Lo Monaco, E.; Tremante, E.; Fruci, D.; Colonna, M.; Giacomini, P. N-linked glycosylation selectively regulates the generic folding of HLA-Cw1. Journal of Biological Chemistry 2008, 283, 16469–16476, doi:10.1074/jbc.M709175200.

43. Baba, E.; Erskine, R.; Boyson, J.E.; Cohen, G.B.; Davis, D.M.; Malik, P.; Mandelboim, O.; Reyburn, H.T.; Strominger, J.L. N-linked carbohydrate on human leukocyte antigen-C and recognition by natural killer cell inhibitory receptors. Human Immunology 2000, 61, 1202–1218, doi:10.1016/s0198-8859(00)00184-1.

44. Ryan, S.O.; Cobb, B.A. Roles for major histocompatibility complex glycosylation in immune function. Semin Immunopathol 2012, 34, 425–441, doi:10.1007/s00281-012-0309-9.

45. Hoek, M.; Demmers, L.C.; Wu, W.; Heck, A.J.R. Allotype-Specific Glycosylation and Cellular Localization of Human Leukocyte Antigen Class I Proteins. J Proteome Res 2021, 20, 4518–4528, doi:10.1021/acs.jproteome.1c00466.

46. Poloski, E.; Oettel, A.; Ehrentraut, S.; Luley, L.; Costa, S.D.; Zenclussen, A.C.; Schumacher, A. JEG-3 Trophoblast Cells Producing Human Chorionic Gonadotropin Promote Conversion of Human CD4+FOXP3-T Cells into CD4+FOXP3+ Regulatory T Cells and Foster T Cell Suppressive Activity. Biology of Reproduction 2016, 94, 106, doi:10.1095/biolreprod.115.135541.

47. Diao, L.H.; Li, G.G.; Zhu, Y.C.; Tu, W.W.; Huang, C.Y.; Lian, R.C.; Chen, X.; Li, Y.Y.; Zhang, T.; Huang, Y.;, et al. Human chorionic gonadotropin potentially affects pregnancy outcome in women with recurrent implantation failure by regulating the homing preference of regulatory T cells. Am J Reprod Immunol 2017, 77, doi:10.1111/aji.12618.

48. Okuyama, M.; Mezawa, H.; Kawai, T.; Urashima, M. Elevated soluble PD-L1 in pregnant women’s serum suppresses the immune reaction. Frontiers in Immunology 2019, 10, 86.

49. Zhang, Y.H.; Aldo, P.; You, Y.; Ding, J.; Kaislasuo, J.; Petersen, J.F.; Lokkegaard, E.; Peng, G.; Paidas, M.J.; Simpson, S. Trophoblast-secreted soluble-PD-L1 modulates macrophage polarization and function. Journal of Leukocyte Biology 2020, 108, 983–998.

50. Warren, J.; Im, M.; Ballesteros, A.; Ha, C.; Moore, T.; Lambert, F.; Lucas, S.; Hinz, B.; Dveksler, G. Activation of latent transforming growth factor-β1, a conserved function for pregnancy-specific beta 1-glycoproteins. Mol. Hum. Reprod. 2018, 24, 602–612.

51. Snyder, S.K.; Wessner, D.H.; Wessells, J.L.; Waterhouse, R.M.; Wahl, L.M.; Zimmermann, W.; Dveksler, G.S. Pregnancy-specific glycoproteins function as immunomodulators by inducing secretion of IL-10, IL-6 and TGF-beta1 by human monocytes. American Journal of Reproductive Immunology 2001, 45, 205–216, doi:10.1111/j.8755-8920.2001.450403.x.

52. Jones, K.; Ballesteros, A.; Mentink-Kane, M.; Warren, J.; Rattila, S.; Malech, H.; Kang, E.; Dveksler, G. PSG9 Stimulates Increase in FoxP3+ Regulatory T-Cells through the TGF-β1 Pathway. PLoS One 2016, 11, e0158050, doi:10.1371/journal.pone.0158050.

53. Huang, X.; Cai, Y.; Ding, M.; Zheng, B.; Sun, H.; Zhou, J. Human chorionic gonadotropin promotes recruitment of regulatory T cells in endometrium by inducing chemokine CCL2. J Reprod Immunol 2020, 137, 102856, doi:10.1016/j.jri.2019.102856.

54. Schumacher, A.; Brachwitz, N.; Sohr, S.; Engeland, K.; Langwisch, S.; Dolaptchieva, M.; Alexander, T.; Taran, A.; Malfertheiner, S.F.; Costa, S.-D. Human chorionic gonadotropin attracts regulatory T cells into the fetal-maternal interface during early human pregnancy. The Journal of Immunology 2009, 182, 5488–5497, doi:10.4049/jimmunol.0803177.

55. Handschuh, K.; Guibourdenche, J.; Tsatsaris, V.; Guesnon, M.; Laurendeau, I.; Evain-Brion, D.; Fournier, T. Human chorionic gonadotropin produced by the invasive trophoblast but not the villous trophoblast promotes cell invasion and is down-regulated by peroxisome proliferator-activated receptor-gamma. Endocrinology 2007, 148, 5011–5019, doi:10.1210/en.2007-0286.

56. Campuzano, M.; Bueno-Sanchez, J.; Agudelo-Jaramillo, B.; Quintana-Castillo, J.C.; Chaouat, G.C.; Maldonado-Estrada, J.G. Glycan expression in chorionic villi from histocultures of women with early-onset preeclampsia: Immunomodulatory effects on peripheral natural killer cells. Journal of Reproductive Immunology 2020, 142, 103212, doi:10.1016/j.jri.2020.103212.

57. Rizzuto, G.; Brooks, J.F.; Tuomivaara, S.T.; McIntyre, T.I.; Ma, S.; Rideaux, D.; Zikherman, J.; Fisher, S.J.; Erlebacher, A. Establishment of fetomaternal tolerance through glycan-mediated B cell suppression. Nature 2022, 603, 497–502, doi:10.1038/s41586-022-04471-0.

58. Abeln, M.; Albers, I.; Peters-Bernard, U.; Flächsig-Schulz, K.; Kats, E.; Kispert, A.; Tomlinson, S.; Gerardy-Schahn, R.; Münster-Kühnel, A.; Weinhold, B. Sialic acid is a critical fetal defense against maternal complement attack. J Clin Invest 2019, 129, 422–436, doi:10.1172/jci99945.

59. Ryan, S.O.; Bonomo, J.A.; Zhao, F.; Cobb, B.A. MHCII glycosylation modulates Bacteroides fragilis carbohydrate antigen presentation. J Exp Med 2011, 208, 1041–1053, doi:10.1084/jem.20100508.

60. Daly, J.; Sarkar, S.; Natoni, A.; Stark, J.C.; Riley, N.M.; Bertozzi, C.R.; Carlsten, M.; O’Dwyer, M.E. Targeting hypersialylation in multiple myeloma represents a novel approach to enhance NK cell-mediated tumor responses. Blood Adv 2022, 6, 3352–3366, doi:10.1182/bloodadvances.2021006805.

61. Chao de la Barca, J.M.; Chabrun, F.; Lefebvre, T.; Roche, O.; Huetz, N.; Blanchet, O.; Legendre, G.; Simard, G.; Reynier, P.; Gascoin, G. A Metabolomic Profiling of Intra-Uterine Growth Restriction in Placenta and Cord Blood Points to an Impairment of Lipid and Energetic Metabolism. Biomedicines 2022, 10, doi:10.3390/biomedicines10061411.

62. Kandarian, F.; Sunga, G.M.; Arango-Saenz, D.; Rossetti, M. A Flow Cytometry-Based Cytotoxicity Assay for the Assessment of Human NK Cell Activity. J Vis Exp 2017, doi:10.3791/56191.

63. Kim, G.G.; Donnenberg, V.S.; Donnenberg, A.D.; Gooding, W.; Whiteside, T.L. A novel multiparametric flow cytometry-based cytotoxicity assay simultaneously immunophenotypes effector cells: comparisons to a 4 h 51Cr-release assay. J Immunol Methods 2007, 325, 51–66, doi:10.1016/j.jim.2007.05.013.

64. Chen, Q.; Pang, P.C.; Cohen, M.E.; Longtine, M.S.; Schust, D.J.; Haslam, S.M.; Blois, S.M.; Dell, A.; Clark, G.F. Evidence for Differential Glycosylation of Trophoblast Cell Types. Mol. Cell. Proteomics 2016, 15, 1857–1866, doi:10.1074/mcp.M115.055798.

65. Hanisch, F.G.; Uhlenbruck, G.; Peter-Katalinic, J.; Egge, H.; Dabrowski, J.; Dabrowski, U. Structures of neutral O-linked polylactosaminoglycans on human skim milk mucins. A novel type of linearly extended poly-N-acetyllactosamine backbones with Gal beta(1-4)GlcNAc beta(1-6) repeating units. J Biol Chem 1989, 264, 872–883.

66. Fukuda, M.; Carlsson, S.R.; Klock, J.C.; Dell, A. Structures of O-linked oligosaccharides isolated from normal granulocytes, chronic myelogenous leukemia cells, and acute myelogenous leukemia cells. J Biol Chem 1986, 261, 12796–12806.

67. Kuan, S.F.; Byrd, J.C.; Basbaum, C.; Kim, Y.S. Inhibition of mucin glycosylation by aryl-N-acetyl-alpha-galactosaminides in human colon cancer cells. J Biol Chem 1989, 264, 19271–19277.

68. Menier, C.; Riteau, B.; Dausset, J.; Carosella, E.D.; Rouas-Freiss, N. HLA-G truncated isoforms can substitute for HLA-G1 in fetal survival. Human immunology 2000, 61, 1118–1125.

69. Carosella, E.D.; Moreau, P.; Le Maoult, J.; Le Discorde, M.; Dausset, J.; Rouas-Freiss, N. HLA-G molecules: from maternal-fetal tolerance to tissue acceptance. Advances in immunology 2003, 81, 199–252.

70. Zhang, L.; Feng, Y.; Zhang, Y.; Sun, X.; Yi, S.; Wang, K.; Ma, X.; Ma, Q.; Ma, F. Defective HLA-G1 glycosylation disrupts Siglec-7-mediated NK cell tolerance at the maternal-fetal interface in recurrent pregnancy loss. Cell Commun Signal 2026, 24, doi:10.1186/s12964-026-02831-1.

71. Rouas-Freiss, N.; Marchal, R.E.; Kirszenbaum, M.; Dausset, J.; Carosella, E.D. The alpha1 domain of HLA-G1 and HLA-G2 inhibits cytotoxicity induced by natural killer cells: is HLA-G the public ligand for natural killer cell inhibitory receptors? Proc Natl Acad Sci U S A 1997, 94, 5249–5254, doi:10.1073/pnas.94.10.5249.

72. Freeze, H.H.; Kranz, C. Endoglycosidase and glycoamidase release of N-linked glycans. Curr Protoc Mol Biol 2010, *Chapter* 17, Unit 17.13A, doi:10.1002/0471142727.mb1713as89.

73. Gray, C. Glycoprotein gonadotropins. Structure and synthesis. Acta Endocrinologica. Supplementum 1988, 288, 20–27.

74. de Medeiros, S.F.; Norman, R.J. Human choriogonadotrophin protein core and sugar branches heterogeneity: basic and clinical insights. Human Reproduction Update 2009, 15, 69–95, doi:10.1093/humupd/dmn036.

75. Cole, L.A. hCG, the wonder of today’s science. Reproductive Biology and Endocrinology 2012, 10, 1–18.

76. Park, G.M.; Lee, S.; Park, B.; Kim, E.; Shin, J.; Cho, K.; Ahn, K. Soluble HLA-G generated by proteolytic shedding inhibits NK-mediated cell lysis. Biochem Biophys Res Commun 2004, 313, 606–611, doi:10.1016/j.bbrc.2003.11.153.

77. Apps, R.; Murphy, S.P.; Fernando, R.; Gardner, L.; Ahad, T.; Moffett, A. Human leucocyte antigen (HLA) expression of primary trophoblast cells and placental cell lines, determined using single antigen beads to characterize allotype specificities of anti-HLA antibodies. Immunology 2009, 127, 26–39, doi:10.1111/j.1365-2567.2008.03019.x.

78. Lee, C.Q.; Gardner, L.; Turco, M.; Zhao, N.; Murray, M.J.; Coleman, N.; Rossant, J.; Hemberger, M.; Moffett, A. What is trophoblast? A combination of criteria define human first-trimester trophoblast. Stem Cell Reports 2016, 6, 257–272.

79. Blaschitz, A.; Weiss, U.; Dohr, G.; Desoye, G. Antibody reaction patterns in first trimester placenta: implications for trophoblast isolation and purity screening. Placenta 2000, 21, 733–741.

80. Mühlhauser, J.; Crescimanno, C.; Kasper, M.; Zaccheo, D.; Castellucci, M. Differentiation of human trophoblast populations involves alterations in cytokeratin patterns. Journal of Histochemistry & Cytochemistry 1995, 43, 579–589.

81. Mirkovic, J.; Elias, K.; Drapkin, R.; Barletta, J.A.; Quade, B.; Hirsch, M.S. GATA3 expression in gestational trophoblastic tissues and tumours. Histopathology 2015, 67, 636–644, doi:10.1111/his.12681.

82. Ma, G.T.; Roth, M.E.; Groskopf, J.C.; Tsai, F.-Y.; Orkin, S.H.; Grosveld, F.; Engel, J.D.; Linzer, D. GATA-2 and GATA-3 regulate trophoblast-specific gene expression in vivo. Development 1997, 124, 907–914.

83. Ralston, A.; Cox, B.J.; Nishioka, N.; Sasaki, H.; Chea, E.; Rugg-Gunn, P.; Guo, G.; Robson, P.; Draper, J.S.; Rossant, J. Gata3 regulates trophoblast development downstream of Tead4 and in parallel to Cdx2. Development 2010, 137, 395–403.

84. Home, P.; Ray, S.; Dutta, D.; Bronshteyn, I.; Larson, M.; Paul, S. GATA3 is selectively expressed in the trophectoderm of peri-implantation embryo and directly regulates Cdx2 gene expression. Journal of Biological Chemistry 2009, 284, 28729–28737.

85. Apps, R.; Sharkey, A.; Gardner, L.; Male, V.; Trotter, M.; Miller, N.; North, R.; Founds, S.; Moffett, A. Genome-wide expression profile of first trimester villous and extravillous human trophoblast cells. Placenta 2011, 32, 33–43.

86. Saben, J.; Zhong, Y.; McKelvey, S.; Dajani, N.K.; Andres, A.; Badger, T.M.; Gomez-Acevedo, H.; Shankar, K. A comprehensive analysis of the human placenta transcriptome. Placenta 2014, 35, 125–131.

87. Jandus, C.; Boligan, K.F.; Chijioke, O.; Liu, H.; Dahlhaus, M.; Démoulins, T.; Schneider, C.; Wehrli, M.; Hunger, R.E.; Baerlocher, G.M.;, et al. Interactions between Siglec-7/9 receptors and ligands influence NK cell-dependent tumor immunosurveillance. J Clin Invest 2014, 124, 1810–1820, doi:10.1172/jci65899.

88. Rillahan, C.D.; Antonopoulos, A.; Lefort, C.T.; Sonon, R.; Azadi, P.; Ley, K.; Dell, A.; Haslam, S.M.; Paulson, J.C. Global metabolic inhibitors of sialyl- and fucosyltransferases remodel the glycome. Nature Chemical Biology 2012, 8, 661–668, doi:10.1038/nchembio.999.

89. Alter, G.; Malenfant, J.M.; Altfeld, M. CD107a as a functional marker for the identification of natural killer cell activity. J Immunol Methods 2004, 294, 15–22, doi:10.1016/j.jim.2004.08.008.

90. De Oliveira, L.G.; Lash, G.E.; Murray-Dunning, C.; Bulmer, J.N.; Innes, B.A.; Searle, R.F.; Sass, N.; Robson, S.C. Role of Interleukin 8 in Uterine Natural Killer Cell Regulation of Extravillous Trophoblast Cell Invasion. Placenta 2010, 31, 595–601, doi:10.1016/j.placenta.2010.04.012.

91. Hu, Y.; Dutz, J.P.; MacCalman, C.D.; Yong, P.; Tan, R.; von Dadelszen, P. Decidual NK cells alter in vitro first trimester extravillous cytotrophoblast migration: a role for IFN-gamma. J. Immunol. 2006, 177, 8522–8530, doi:10.4049/jimmunol.177.12.8522.

92. Lash, G.E.; Otun, H.A.; Innes, B.A.; Kirkley, M.; De Oliveira, L.; Searle, R.F.; Robson, S.C.; Bulmer, J.N. Interferon-gamma inhibits extravillous trophoblast cell invasion by a mechanism that involves both changes in apoptosis and protease levels. FASEB J. 2006, 20, 2512–2518, doi:10.1096/fj.06-6616com.

93. Zhou, Y.; Fu, B.; Xu, X.; Zhang, J.; Tong, X.; Wang, Y.; Dong, Z.; Zhang, X.; Shen, N.; Zhai, Y. PBX1 expression in uterine natural killer cells drives fetal growth. Science Translational Medicine 2020, 12, eaax1798.

94. Ma, Y.; Yang, Q.; Fan, M.; Zhang, L.; Gu, Y.; Jia, W.; Li, Z.; Wang, F.; Li, Y.X.; Wang, J.;, et al. Placental endovascular extravillous trophoblasts (enEVTs) educate maternal T-cell differentiation along the maternal-placental circulation. Cell Proliferation 2020, 53, e12802, doi:10.1111/cpr.12802.

95. Segerer, S.E.; Müller, N.; Van Den Brandt, J.; Kapp, M.; Dietl, J.; Reichardt, H.M.; Rieger, L.; Kämmerer, U. Impact of female sex hormones on the maturation and function of human dendritic cells. American Journal of Reproductive Immunology 2009, 62, 165–173.

96. Mkhikian, H.; Mortales, C.L.; Zhou, R.W.; Khachikyan, K.; Wu, G.; Haslam, S.M.; Kavarian, P.; Dell, A.; Demetriou, M. Golgi self-correction generates bioequivalent glycans to preserve cellular homeostasis. Elife 2016, 5, doi:10.7554/eLife.14814.

97. Takamatsu, S.; Antonopoulos, A.; Ohtsubo, K.; Ditto, D.; Chiba, Y.; Le, D.T.; Morris, H.R.; Haslam, S.M.; Dell, A.; Marth, J.D.;, et al. Physiological and glycomic characterization of N-acetylglucosaminyltransferase-IVa and -IVb double deficient mice. Glycobiology 2010, 20, 485–497, doi:10.1093/glycob/cwp200.

98. Deng, Q.; Liu, X.; Yang, Z.; Xie, L. Expression of N-Acetylglucosaminyltransferase III Promotes Trophoblast Invasion and Migration in Early Human Placenta. Reproductive Sciences 2019, 26, 1373–1381, doi:10.1177/1933719118765967.

99. Endo, T.; Nishimura, R.; Kawano, T.; Mochizuki, M.; Kobata, A. Structural differences found in the asparagine-linked sugar chains of human chorionic gonadotropins purified from the urine of patients with invasive mole and with choriocarcinoma. Cancer Res 1987, 47, 5242–5245.

100. Blithe, D.L. N-linked oligosaccharides on free alpha interfere with its ability to combine with human chorionic gonadotropin-beta subunit. Journal of Biological Chemistry 1990, 265, 21951–21956.

101. Cao, L.; Wang, Y.; Ruan, L.; Chen, H.; Yang, S.; Xu, J.; Ma, H.; Luo, X.; Xie, Y.; Chen, E.;, et al. Protein N-glycosylation supports extravillous trophoblast cell lineage development in hypoxia microenvironment. Life Sciences 2026, 124518, doi:doi.org/10.1016/j.lfs.2026.124518.

102. Zhang, L.; Feng, Y.; Wu, P.; Chen, L.; Jiang, N.; Ma, X.; Ma, Q.; Lu, H.J.; Xiao, X.; Ma, F. Deficient extravillous trophoblast invasion caused by impaired sialylation-Siglec-7 interaction contributes to recurrent pregnancy loss. Cell Death Dis 2026, 17, doi:10.1038/s41419-026-08503-9.

103. Huang, J.; Feng, L.; Huang, J.; Zhang, G.; Liao, S. Unveiling sialoglycans’ immune mastery in pregnancy and their intersection with tumor biology. Front Immunol 2024, 15, 1479181, doi:10.3389/fimmu.2024.1479181.

