## Supplementary Tables for "Extravillous trophoblast model shows generation of bioequivalent N-glycans can maintain immunological protection against natural killer cell cytotoxicity"

**Supplementmentary Table 1. Inhibitors for different N-glycan processing enzymes.**

| **Inhibitor** | **Structure** | **Targeting enzyme** | **Working conc.** |
| --- | --- | --- | --- |
| Kifunensine (KIF) | 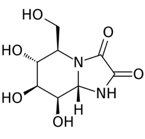 | α-mannosidase I resident in both ER and Golgi | 10 μg/mL |
| Swainsonine (SWA) | 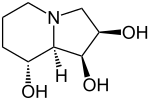 | α-mannosidase II resident in Golgi apparatus | 10 μg/mL |
| 3F_AX_-Peracetyl Neu5Ac (3FN) | 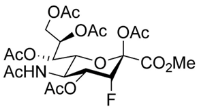 | Sialyltransferase | 300 µM |
| Benzyl-α-GalNAc (BAG) | 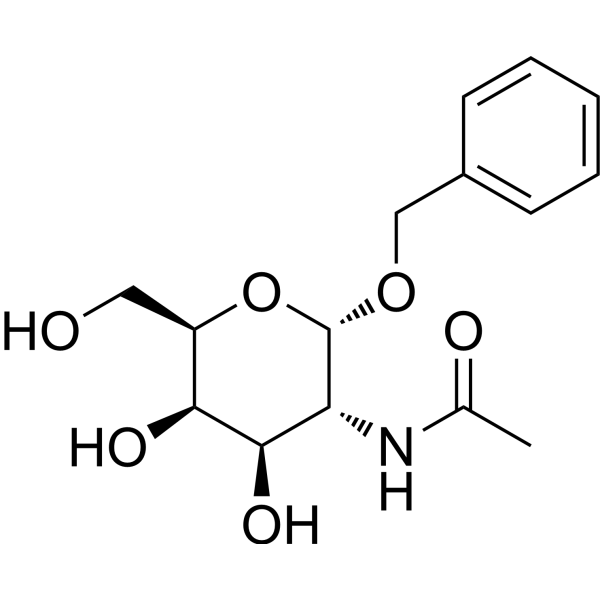 | UDP-Gal:GalNAcα peptide β-1,3-galactosyltransferases (C1Gal-Ts) | 2 mM |

**Supplementmentary Table 2. Representative plant lectins for cell surface glycan analysis.**

| **Lectin** | **Glycan-binding specificity** | **References** |
| --- | --- | --- |
| *Canavalia ensiformis* agglutinin (ConA) | Mannose, glucose | [1] |
| *Lens culinaris* lectin (LCA) | Core α-1,6-linked fucose |  |
| *Lycopersicon esculentum* lectin (LEL) | Polylactosamine chains |  |
| *Phaseolus vulgaris* erythronagglutinin (PHA-E) | Bisecting β-1,4-GlcNAc branch |  |
| *Phaseolus vulgaris* leukoaggulutinin (PHA-L) | β-1,6-GlcNAc branch |  |
| *Sambucus nigra* agglutinin (SNA) | α-2,6-linked sialic acid |  |

Supplementmentary Table 3. Characteristics of different anti-HLA-G mAbs used in this study.

| **Clone** | **Forms of**  **HLA-G detected** | **Conformation of HLA-G detected** | **Immuno-**  **reacted with** | **Conjugate** | **Applications in this study** |
| --- | --- | --- | --- | --- | --- |
| G233 | Transmembrane and soluble | Conformation-dependant | Native  HLA-G1 [2] | None | Immuno-precipitation |
| MEM-G/9 | Transmembrane | Conformation-dependant | Native HLA-G1 and HLA-G3 [2,3] | APC, PE or FITC | Flow cytometry |
| 4H84 | Transmembrane and soluble | Denatured/  unfolded | Denatured HLA-G1, -G2, -G3 and HLA-G4 [2] | AF488 or HRP | Flow cytometry and Western blotting |

**Note:** AF488, Alexa Fluor® 488, APC, allophycocyanin; FITC, fluorescein Isothiocyanate; HRP, horseradish peroxidase; PE, phycoerythrin;

**References**

1.	Bojar, D.; Meche, L.; Meng, G.; Eng, W.; Smith, D.F.; Cummings, R.D.; Mahal, L.K. A Useful Guide to Lectin Binding: Machine-Learning Directed Annotation of 57 Unique Lectin Specificities. *ACS Chem Biol* **2022**, *17*, 2993-3012, doi:10.1021/acschembio.1c00689.

2.	Furukawa, A.; Meguro, M.; Yamazaki, R.; Watanabe, H.; Takahashi, A.; Kuroki, K.; Maenaka, K. Evaluation of the reactivity and receptor competition of HLA-G isoforms toward available antibodies: implications of structural characteristics of HLA-G isoforms. *International journal of molecular sciences* **2019**, *20*, 5947.

3.	Zhao, L.; Teklemariam, T.; Hantash, B. Reassessment of HLA‐G isoform specificity of MEM‐G/9 and 4H84 monoclonal antibodies. *Tissue antigens* **2012**, *80*, 231-238.
